# Skeleton-forming responses of reef-building corals under ocean acidification

**DOI:** 10.1101/2024.05.25.595876

**Authors:** Yixin Li, Hongwei Zhao, Yunpeng Zhao, Xin Liao, J.-Y. Chen, Chunpeng He, Zuhong Lu

## Abstract

Ocean acidification is increasing in frequency and is considered one of the most important causes of severe damage to global coral reefs. Therefore, there is an urgent need to study the impact of acid stress on the growth patterns of major reef-building corals. Here, we studied the skeleton forming strategies of four widely distributed coral species in a simulated acidified habitat with a pH of 7.6–7.8. We reconstructed and visualized the skeleton building process, quantified elemental calcium loss, and determined gene expression changes. The results suggest that different reef-building corals have diverse growing strategies in acidified seawater. A unique ‘cavity-like’ forming process starts from the inside of the skeletons of *Acropora muricata*, which sacrifices skeleton density to protect its polyp-canal system. The forming patterns in *Pocillopora damicornis*, *Montipora capricornis*, and *M. foliosa* were characterized by ‘osteoporosis’, exhibiting disordered skeletal structures, insufficient synthesis of adhesion proteins, and low bone mass, correspondingly. In addition, we found that skeletal areas near coral polyps suffered less and had later acidified damage than other skeletal areas in the colony. These results help to understand the skeleton-forming strategies of several major coral species under acid stress, thereby laying a foundation for coral reef protection and restoration under increasing ocean acidification.

## Introduction

Scleractianian reef-building corals appeared on earth 250 Myr (million years) ago and have experienced unprecedentedly sudden changes in oceanic pH in the last two centuries^1^. The anthropogenic production of carbon dioxide (CO_2_) has decreased the pH by an average of 0.1 since the Industrial Revolution (about 200 years ago), with even higher changes occurring in heavily polluted offshore areas^2^. In the previous 20 Myr, the oceanic pH changed by less than ± 0.3. According to the forecast from Intergovernmental Panel on Climate Change (IPCC), the global average seawater pH will continue to drop to 7.8–7.9 by 2100 (current pH is 8.1 ± 0.1), potentially posing devastating effects on organisms with calcium carbonate skeletons, such as corals^3^.

Recent research suggests that the acidification of global oceans leads to a reduction in the diversity, biomass, and trophic complexity of benthic marine communities, resulting in significant negative impacts on certain marine species with potential ecosystem-level consequences, posing a threat to the biodiversity and ecosystem function of coral reef ecosystems^4–6^. It is anticipated that the net calcification of reef-building corals will decline as ocean acidity increases, potentially leading to net dissolution by the end of this century^7,8^. Furthermore, habitats with lower pH can disrupt calcification efficiency, coral-Symbiodiniaceae holobiont functioning, self-healing ability, and larval attachment due to lack of natural resistance mechanisms in some reef-building corals^9–13^. However, evidence suggests that reef-building corals may have inherent resilience against extinction events caused by ocean acidification throughout their life history based on past large-scale extinctions accompanied by severe ocean acidification^14^.

In this study, we investigated the impact of an acidified environment on the growth process of reef-building corals by examining the growth and mineralization responses of four widely distributed coral species (*Acropora muricata*, *Montipora capricornis*, *M. foliosa*, and *Pocillopora damicornis*) under acid stress. We simulated an ocean acidification habitat for these coral samples with pH values fluctuating rhythmically between 7.8 during the day and 7.6 at night (control conditions maintained pH values between 8.2 during the day and 8.0 at night). The response of coral polyps in a colony and their calcium carbonate skeletons to external influences was jointly regulated by the polyp-canal system^15–17^. We collected samples from these four corals in our simulation system at Days 0, 3, 6, 9, and 30 for high-resolution micro-computed tomography (micro-CT), scanning electron microscopy with energy dispersive spectrometry (SEM-EDS), and transcriptome sequencing (RNA-seq) analyses. Reconstructions of the polyp-canal system were used to visualize the growth process in coral colonies and quantify skeleton loss during the 30-day acid stress exposure. Elemental ratios in the coral skeletons were analyzed to determine calcium loss/deposition trends while gene expression changes in the coral skeletome^18–19^ revealed impacts on biomineralization^20^.

In summary, we present insights into skeleton-forming strategies of these representative coral species that highlight effects of excessive CO_2_ on coral growth; providing a theoretical basis for protecting reef-building corals and restoring acid-affected reefs under global ocean acidification.

## Results

### Erosion process in the skeletal system

We reconstructed the polyp-canal systems^21^ in each coral colony of *A. muricata*, *M. foliosa*, *M. capricornis*, and *P. damicornis* samples from Day 0 to Day 30, visualizing their erosion process under acid stress (Figures 1, S1–21).

**Figure 1.**
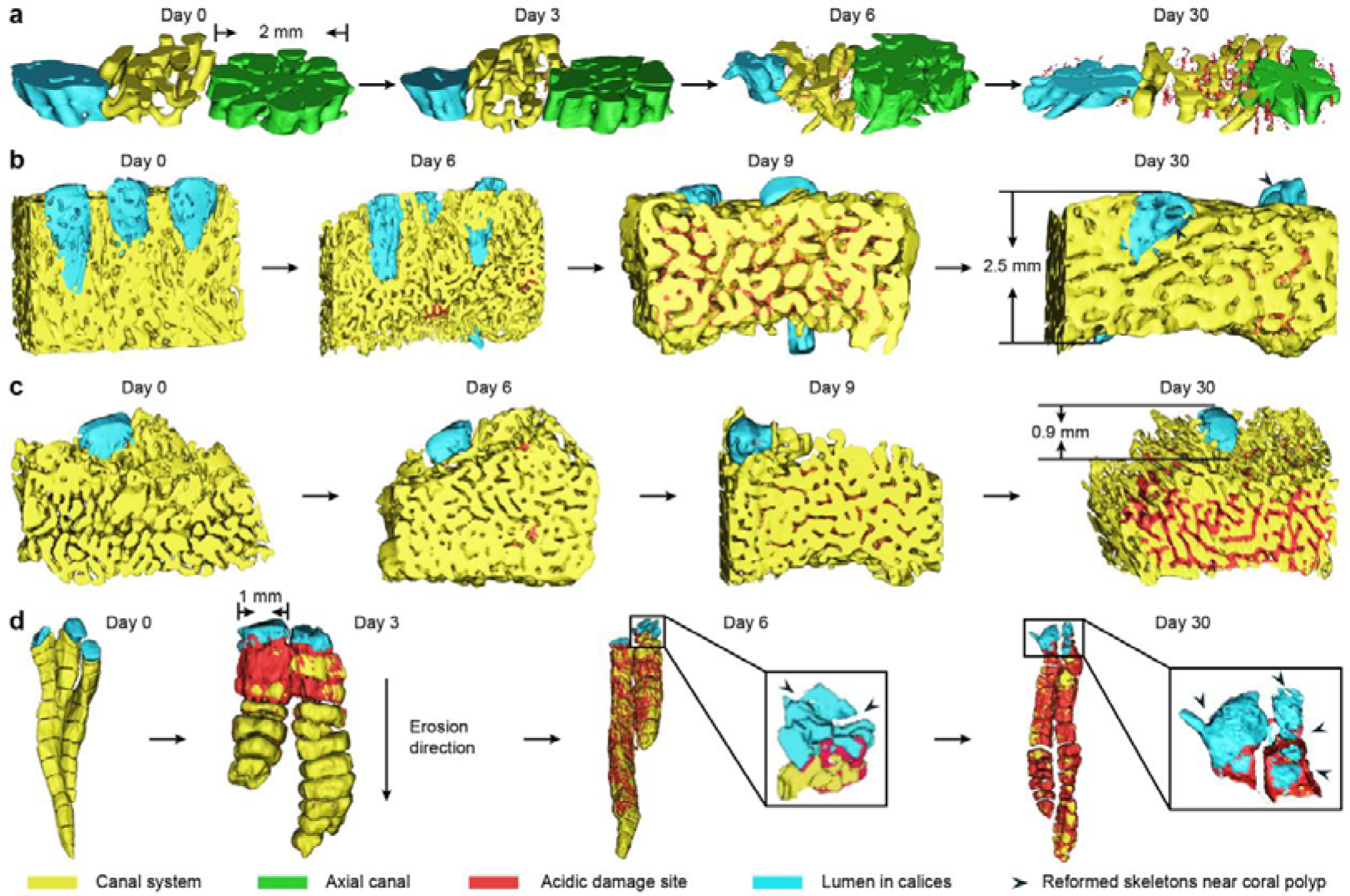
3D reconstructions of polyp-canal systems visualized the corrosion process in reef-building corals under acidic stress. a) In *A. muricata*, corrosion occurred inside coral skeletons near canal system at Day 3, and was firstly found in calices at Day 6. Till Day 30, the ‘cavity-like’ acidic damage sites had spread inside skeletons of the colony. b) In *M. capricornis*, corrosion occurred on the surface of coral skeletons at Day 6. The acidic damage sites spread on skeletons surrounding the canal system at Day 9, while they were also found in calices near polyps. However, most acidic damage sites were covered by newly formed skeletons at Day 30, and some of them could still be noticed over the canal system away from polyps. c) In *M. foliosa*, corrosion occurred on the surface of coral skeletons at Day 6 and could be found in calices at Day 9. The acidic damage sites kept spreading at Day 30, while the skeletons near coral polyps suffered less damages than those away from polyps. d) In *P. damicornis*, corrosion occurred over skeletons near the surface of coral colony at Day 3, and spread deep into the colony along the walls and dissepiments around inter-septal spaces. Reformed skeletons occurred in calices at Day 9, covering the acidic damage sites, while irregular reformed skeletons were found in several lumens of the calices, squeezing the lebensraum of coral polyps.

In *A. muricata*, the initial occurrence of acidic damage sites was observed within the skeletons near the canal system on Day 3 (Figures 1a, S2). Subsequently, there was a continuous increase in both the quantity and volume of these pore-like sites in the following days. By Day 6, acidic damage sites were also present in axial skeletons and calice walls near coral polyps (Figure S3), leading to internal corrosion and surface erosion across all skeletons within the A. muricata colony (Figures S4, S5). The adjacent small pore-like acid sites gradually merged to form larger porous sites or fractures within the skeletal structure (Figure S21a).

In the other three coral species, the erosion processes started at the surfaces of coral skeletons. In *M. capricornis* and *M. foliosa*, acid corrosion areas occurred on the surface of the skeletons at Day 6, which increased the volume of the canal system (Figures 1b, 1c, S8, S13). We could not find any corrosion areas over the calice walls until Day 9, and the corrosion areas had already spread over most skeletons around the canal system (Figures S9, S14). Fewer corrosion areas remained in the *M. capricornis* colony at Day 30, and the structure of the polyp-canal system was similar to that prior to acid stress (Figures S10, S21b). A small amount of irregular reformed skeletons caused tiny gaps in the original canal system and lumen in the calices of the coral colony (Figure 1b). However, the corrosion areas surrounding the canal system continued to expand until Day 30 in *M. foliosa* (Figure 1c), while the corrosive impact near coral polyps was notably less pronounced during this period (Figure S21c). The polyp-canal system in *M. foliosa* suffered serious damage under acid stress, which had implications for coral growth (Figure S15).

Corrosion areas first occurred in *P. damicornis* at Day 3 over the walls of coral calices and dissepiments near the surface of the coral colony (Figures 1d, S17). From Day 3 to Day 6, the corrosion areas spread inside the colony along the coenosteums around inter-septal spaces in each calice (Figure S21d). The corrosion areas near coral polyps became smaller and the appearance of reformed skeletons fragmented the lumen in the calices (Figure S18). At Day 30, the network of inter-septal spaces in the *P. damicornis* colony had become seriously damaged, corrosion areas covered most of the internal skeletons, and some dissepiments and coenosteums were broken off, causing adjacent inter-septal spaces to merge into larger ones (Figures 1d, S20). In contrast, more reformed skeletons occurred in several lumens of the calices, further squeezing the lebensraum of these polyps (Figures 1d, S21d).

In conclusion, under acid stress, acid corrosion areas initiated inside the coral skeletons in *A. muricata* and started from their surfaces in *M. capricornis*, *M. foliosa*, and *P. damicornis*. Among these four species, the polyp-canal systems of *A. muricata* and *M. capricornis* were less affected during the 30-day ocean acidification simulation. In addition, we found that skeletons close to coral polyps were less affected by acid stress, with corrosion appearing later in those regions than in other regions of the colony, except in *P. damicornis*.

### Skeleton and calcium loss

We quantified skeletal loss during the 30-day acidification experiment by analyzing the skeleton to void ratio (Figure S22a, Table S1). We randomly selected nine 1 mm × 1 mm × 1 mm cuboid areas in each sample and reconstructed their polyp-canal systems. The trends in the average skeleton to void ratio in each coral species all decreased with experiment time (Figure 2). The average volume of canals, as a percentage of the total coral volume in *A. muricata* increased from 39.4% to 49.7%, which meant that nearly 17.0% (10.3%/60.6%) of the calcareous skeleton in this colony was lost during the acid stress experiment (Figure 2a). In *M. foliosa*, the canal volume as a percentage of total volume increased from 67.0% to 73.0%, while about 18.2% (6.0%/33.0%) of the coral skeleton was corroded. We found that the vast majority of skeleton loss occurred in the first nine days, with significantly lower losses after Day 9 (Figure 2b). In *M. capricornis*, the canal volume ratio grew from 54.1% to 64.7% between Days 0 and 9 and decreased to 61.3% at Day 30. The volume of coral skeleton increased abnormally between Days 9 and 30, which meant that more skeleton was formed than was eroded during the experiment. The skeleton loss was 23.1% (10.6%/45.9%) at Day 9 and decreased to nearly 15.7% (7.2%/45.9%) at Day 30 (Figure 2c). As in *P. damicornis*, the canal volume ratio kept increasing, from 38.8% to 50.8%, and the approximately uniform skeleton loss was 19.6% (12%/61.2%) in total (Figure 2d).

**Figure 2.**
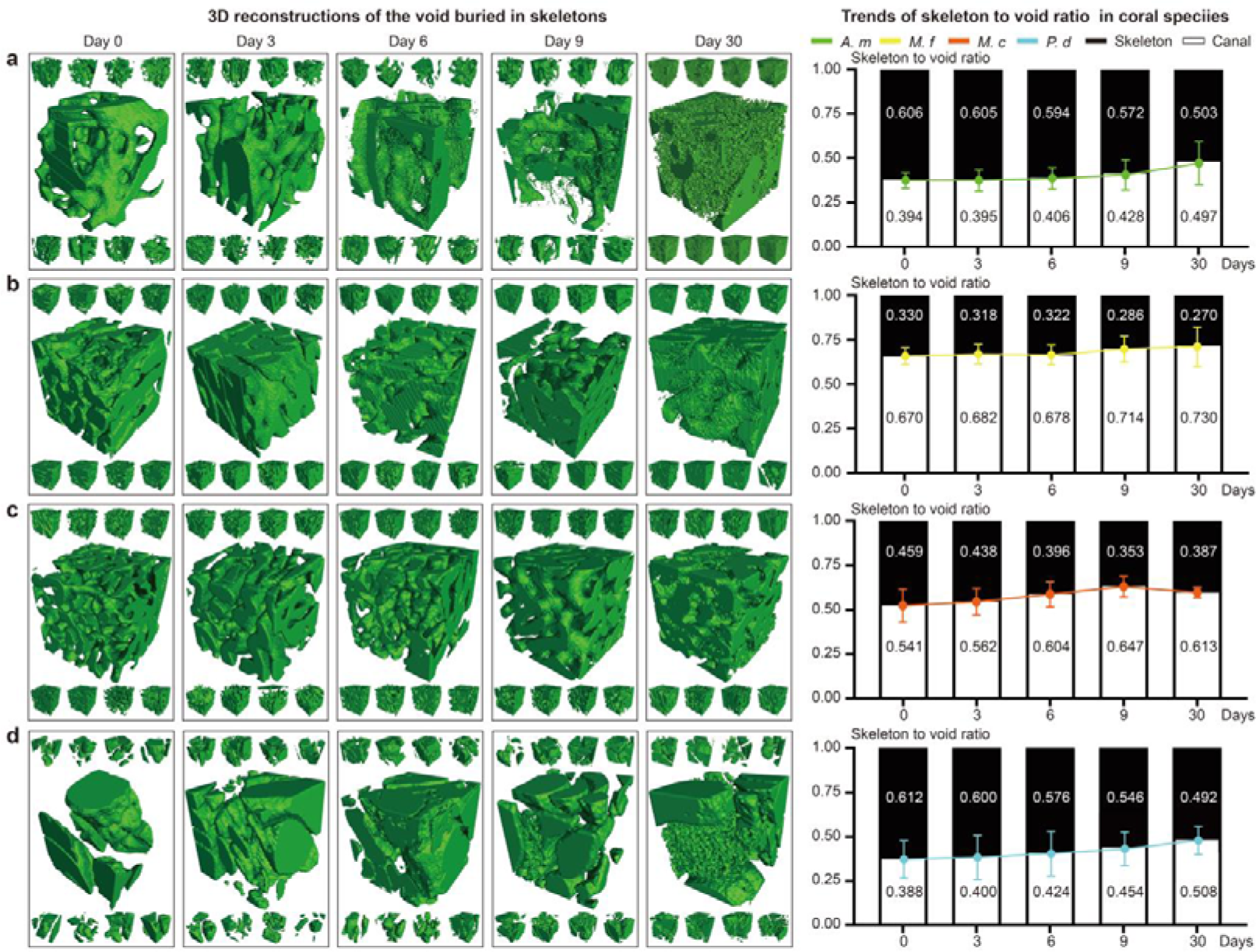
Skeleton to void ratio of coral colony revealed the skeleton loss under acidic stress. The green areas in a-d are the 3D reconstructions of void, which are buried in the skeletons. a) The volume of skeletal matter as a percentage of total coral volume in *A. muricata* colony decreased from 60.6% at Day 0 to 50.3% at Day 30. b) The volume of skeletal matter as a percentage of total coral volume in *M. foliosa* colony decreased from 33.0% at Day 0 to 27.0% at Day 30. c) The volume of skeletal matter as a percentage of total coral volume in *M. capricornis* colony decreased from 45.9% at Day 0 to 35.3% at Day 9, and rebounded to 38.7% at Day 30. d) The volume of skeletal matter as a percentage of total coral volume in *P. damicornis* colony decreased from 61.2% at Day 0 to 49.2% at Day 30.

Calcium (Ca), Oxygen (O), and Carbon (C) are the main elements present in the corallite of reef-building corals. We used SEM-EDS to obtain the variations in the element atomic ratios of the four coral species undergoing acid stress for 30 days (Figures 3a-d, S22b, Table S2). For all four coral species, the atomic ratio of Ca decreased between Days 0 to 30 (Figures 3e, S23-26). Due to the loss of Ca^2+^, the atomic ratio of Ca decreased between Days 0 and 30, with rates of decrease of 5.40% in *A. muricata*, 2.26% in *M. foliosa*, 1.01% in *M. capricornis*, and 5.94% in *P. damicornis* (Figure 3, Table S3). Against the background of these decreasing trends, the Ca atomic ratio remained constant between Days 0 and 3 in *M. foliosa* and *M. capricornis*, and appeared to slightly rebound at Day 6 in *M. foliosa* (from 25.11% to 25.74%) and at Day 30 in *M. capricornis* (from 24.24% to 25.74%).

**Figure 3.**
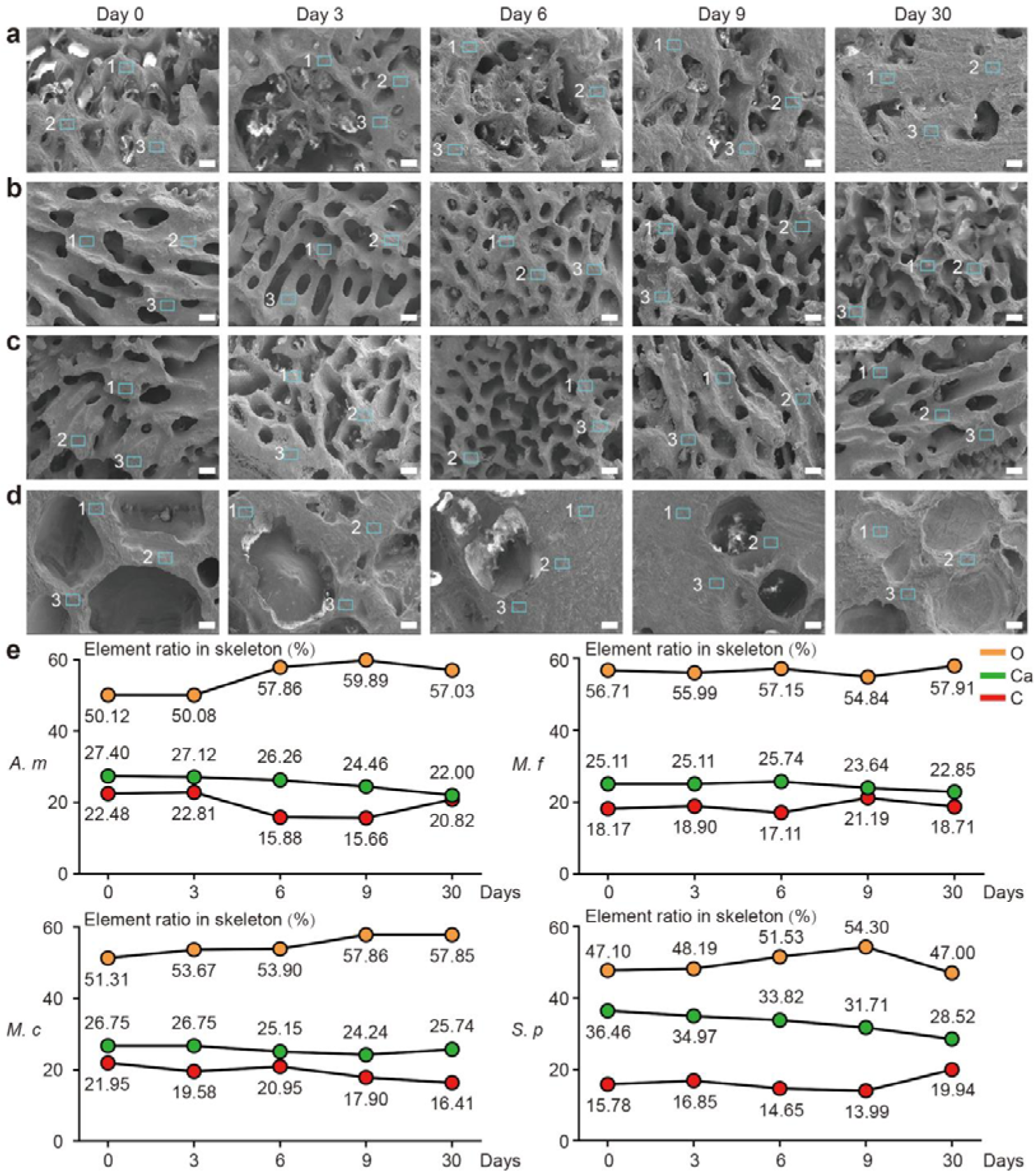
Changes of skeleton element ratio revealed the Ca^2+^ loss in coral colony under acidic stress. a-d) SEM figures of **a** *A. muricata*; **b** *M. foliosa*; **c** *M. capricornis*; and **d** *P. damicornis* from Day 0 to Day 30, and the signals 1, 2, and 3 in each figure are the sampling sites foe EDS test. e) The atomic ratio of Ca decreased 5.40% in *A. muricata* (*A.m*), 2.26% in *M. foliosa* (*M.f*), 1.01% in *M. capricornis* (*M.c*), and 5.94% in *P. damicornis* (*M.d*). Scale bar: 0.2 mm.

### Gene expression changes in the skeletome

The RNA-seq tests showed that the expressions of most genes in these four coral species were affected by ocean acidification at Day 3, which we believe to be the stress condition of these coral-symbiodinium holobionts. The expression of most genes returned to a steady state at Day 9; however, the skeletome in the coral polyps still suffered significant effects (Figure 4). Among the proteins involved in coral skeleton formation and calcium transportation, the following five categories suffered most from acidification (Figures S27, Table S3): (1) Transporting proteins like plasma membrane calcium-transporting ATPase (PMC-t ATPase), plasma membrane calcium ATPase (PMC ATPase), and solute carrier (SC). The ATPases are involved in calcium transportation through the membranes of entodermic and calicoblastic cells, while SC helps to transport bicarbonate. (2) Carbonic anhydrase (CA). These are enzymes for the interconversion of carbon dioxide and bicarbonate that can regulate their balance to affect the pH of coral cells. (3) Acid-rich proteins (ARPs), like skeletal aspartic acid-rich protein (SAARP), acidic skeletal organic matrix protein (ASOMP), secreted acidic protein (SAP), and aspartic and glutamic acid-rich protein (AGARP). These ARPs can deposit calcium carbonate from seawater directly for coral skeleton formation. (4) Skeletal organic matrix proteins (SOMPs), which control coral skeleton formation through bio-mineralization. (5) Adhesion proteins like galaxin and collagen alpha-6(VI) chain-like (α-C6), which cement the calcium carbonate crystals to each other and to coral skeletons.

**Figure 4.**
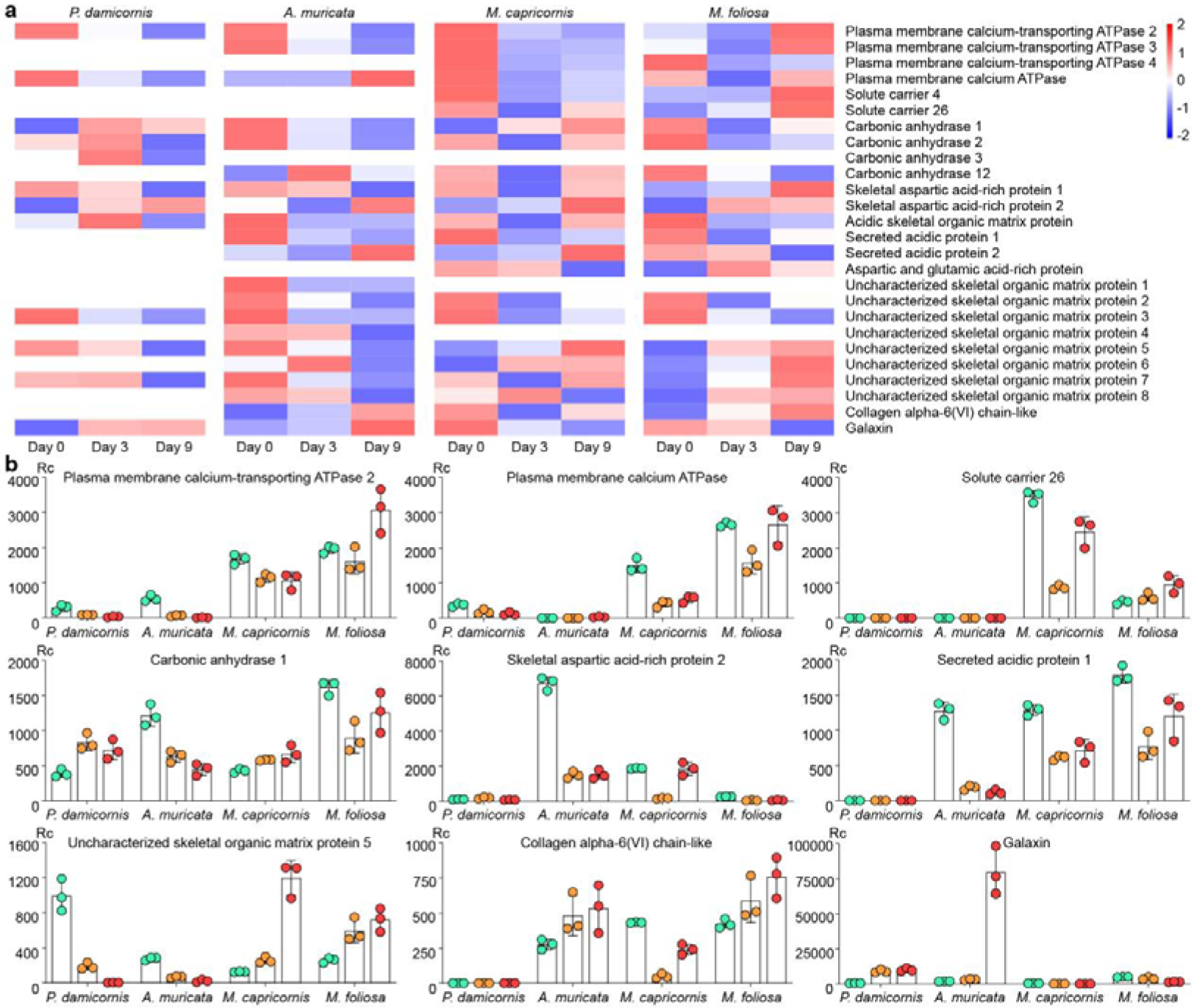
Gene expression changes of skeletomes in the four reef-building corals under acidic stress. (a) The disparity in gene expression of skeletomes at Day 0, Day 3, and Day 9 was shown with a heat map. There were three replicate per time point per species, and the quantities shown in the figure are the average values of the three replicate (detailed data can be found in Table S3). (b) Gene expression changes of nine main proteins. Rc means read count.

In the polyps of *P. damicornis*, nearly half of the skeletomes mentioned above were almost unaffected by acid stress, including SC, SAP, AGARP, SOMP-1/2/4/6/8, and α-C6. The expression of ATPases decreased continuously from Days 0 to 3 and Days 3 to 9. The expressions of CA and ASOMP increased from Days 0 to 3 and decreased from Days 3 to 9. The expression of SAARP-1 decreased while that of SAARP-2 increased from Days 0 to 9. The expressions of SOMP-3/5 decreased continuously from Days 0 to 9, while the expression of SOMP-7 increased from Days 0 to 3 and decreased from Days 3 to 9. The expression of galaxin increased continuously from Days 0 to 9 (Figure 4a). These findings suggest that the polyps of *P. damicornis* exhibits potential for adaptation to ocean acidification, as the expression of approximately half of its skeletome genes remains largely unaffected by acid stress.. The ability to transport calcium through cells was reduced at Day 3, while the ability to adjust pH, and the sedimentation/adhesion of calcium carbonate, were significantly improved. At Day 9, most of the biomineralization abilities were significantly inhibited, except for continuous increases in the adhesion of calcium carbonate (Figure 4b).

In *A. muricata*, the expression of SC was almost unaffected by acid stress. The expressions of ATPases, CA, ARPs and SOMPs decreased continuously from Days 0 to 9. The expressions of α-C6 and galaxin increased continuously from both Days 0 to 3 and Days 3 to 9 (Figure 4a). The acid stress tolerance of *A. muricata* seems weaker than that of *P. damicornis*, and most of its biomineralization processes were significantly inhibited by acidification. However, the secretion of adhesion proteins was greatly increased, which potentially promoted binding among the calcium carbonate crystals that maintain the toughness of the coral colony despite continuous losses of calcium and skeleton (Figure 4b). This pattern may have mitigated the effects of the retardation in skeleton formation.

In *M. capricornis*, the expressions of ATPases and α-C6 decreased from Days 0 to 3 and rebounded slightly from Days 3 to 9. The expressions of SC, CA, SAARP, ASOMP and SAP decreased from Days 0 to 3 and sharply rebounded from Days 3 to 9. The expressions of AGARP and galaxin decreased continuously from Days 0 to 9. The expressions of SOMPs showed a downward trend at Day 3 but significantly rebounded at Day 9 (Figure 4a). The biomineralization ability of *M. capricornis* was significantly inhibited at Day 3 but kept intensifying until Day 9. Despite the compensatory increase in biomineralization processes to counteract skeleton and calcium losses under acid stress, continuous reductions in the secretion of adhesion proteins may render coral colonies more fragile. (Figure 4b).

In *M. foliosa*, the expressions of ATPase and SC decreased from Days 0 to 3 and sharply rebounded from Days 3 to 9. The expressions of CA, ARPs and galaxin showed a downward trend from Days 0 to 9. The expressions of SOMP-2/3 decreased while those of SOMP-5/6/7/8 increased from Days 0 to 9. The expression of α-C6 increased continuously from Days 0 to 9 (Figure 4a). In the *M. foliosa* colony, the abilities to precipitate calcium carbonate from seawater and adjust pH were inhibited under acid stress, while the oriented precipitation of calcium carbonate crystals and the transmembrane transport of calcium and bicarbonate were promoted (Figure 4b).

## Discussion

### Survival strategies of investigated four corals

This work revealed the survival strategies of *A. muricata*, *M. foliosa*, *M. capricornis*, and *P. damicornis* using micro-CT, SEM-EDS, and RNA-seq tests. The ‘cavity-like’ erosion pattern observed in *A. muricata* is different from those of the other three species. Corrosion pores first occurred inside the coral skeletons, which gradually extended to the central area from the near-surface area. During this process, there were no obvious cracks or channels on the surface of the coral skeleton and the polyp-canal system regulating coral growth was barely affected (Figures 1a, S21a). The number and volume of corrosion pores kept increasing in the skeletons under acid stress, while the skeleton to void ratio in the coral colony and the calcium element ratio in the skeletons kept decreasing, and the gene expression of most skeletomes in the coral polyps involved in skeleton formation also decreased significantly (Figures 2a, 3a,e, 4). However, since the polyp-canal system is the basis for the sustainable growth of reef-building corals^17^, this survival strategy—of sacrificing internal structures and calcium inside the skeletons to protect the basic canal network—helps *A. muricata* maintain its growth patterns. Furthermore, the upregulation of galaxin and α-C6 gene expressions may contribute to the maintenance of skeletal toughness and rigidity in spite of the presence of internal cavities, thereby enhancing the protective capacity of the coral colony. (Table S3).

The erosion patterns of the other three coral species are similar to those observed in osteoporosis^23^. They are reflected in the insufficient synthesis of adhesion proteins in *M. capricornis*, which may lead to a decrease in skeletal toughness (Figure 4). The low bone mass and calcium loss in *M. foliosa* decrease the quality and hardness of the coral colony (Figures 2, 3). The microstructure of the coral skeletons is destroyed and become disordered, which may affect the mechanical strength of *P. damicornis* (Figures 1d, S21d). Coral polyps form new skeletons over surrounding corrosion areas through biomineralization, like what osteoblasts do^24^.

The self-protection of *M. capricornis* under acid stress is mainly achieved via its high rate of skeleton formation. The increases in CA, SAARP, ASOMP, SAP, and some SOMPs from Days 3 to 9 facilitated the formation of new coral skeletons, potentially alleviating the impact of skeleton and calcium losses on the coral colonies. (Figures 1b, S21b). This strategy also protects the polyp-canal system in *M. capricornis* and maintains the sustained growth of a coral colony under acid stress (Figures 2, 3). However, the decreasing gene expressions of galaxin and α-C6 affect the toughness of coral skeletons. The erosion pattern resembling osteoporosis renders *M. capricornis* colonies more susceptible to physical damage under the influence of acidification. (Table S3).

Despite the rapid skeleton and calcium losses observed in *M. foliosa* colonies due to acidification damage from Days 6 to 9, there was a significant increase in the expressions of ATPases, SC, SAARP, and some SOMPs by Day 9, which appeared to effectively mitigate further skeleton loss for the remainder of the experiment. (Figures 2-4). This phenomenon was mostly pronounced over or near the walls of calices, while other skeletons in the *M. foliosa* colony still suffering acidic damage (Figures 1c, S21c).

We found that *P. damicornis* polyps have better acid stress tolerance, as the expressions of most genes are barely affected by the acidic habitat (Figure 4). Only half of the genes related to skeleton formation suffered from the erosion process at Day 9, which slightly reduced the biomineralization of the coral polyps (Table S3). Despite this, some of the dissepiments and coenosteums around inter-septal spaces became thinner or even fractured due to the skeleton and calcium losses. The thick and dense calice walls at the surface of the coral colony, where polyps live, were less affected, as new skeletons were formed in the calices that filled part of the corrosion areas over the calice walls. However, several new skeletons were not formed along the original patterns, which separated the lumen in calices into fragmented cubes, reducing the living spaces for polyps (Figures 1d, S21d). Since the coenosarcus between polyps are more likely to break down in an acidic habitat^25^, these irregularly formed skeletons will further aggravate the bail-out of *P. damicornis* polyps. These inducements potentially lead to the disorder and destruction of skeletal structures and affect the mechanical strength of a coral colony.

### Coral polyps protect their surrounding skeletons under acid stress

We found that, in *A. muricata*, *M. capricornis*, and *M. foliosa*, skeletons near coral polyps suffered acidic damage later and to a lesser degree than other skeletons in the same colony. As in *P. damicornis*, although the corrosion areas occurred first over calice walls at the surface of the coral colony around polyps, most of these areas were filled with newly formed skeletons during long-term acid stress, while other skeletons inside the colony away from polyps suffered serious damage (Figure 1). This reveals that, in an acidic habitat, coral polyps with a biomineralization ability can protect the surrounding skeletons by forming new ones. Other skeletons farther away from coral polyps were more severely eroded, which many indicate that the transport of calcium, SOM, calcification fluid, and other skeleton-forming materials via the gastrovascular canals was inhibited by acid stress (Figure S21).

In addition, we also found that although the expressions of various genes related to skeletomes increased in both *M. capricornis* and *M. foliosa* at Day 9, thereby improving their biomineralization ability, polyps in *M. capricornis* protected their skeletons better than those in *M. foliosa* (Figure 4). We believe that this difference is mainly due to the difference between calice volume and intensity. The calices in *M. capricornis* are larger and more densely distributed and border most of the skeletons in the colony, while the calices in *M. foliosa* are much smaller and can only protect nearby skeletons (Figure S11-20).

## Materials and Methods

### Sample collection

The *A. muricata*, *M. capricornis*, *M. foliosa*, and *P. damicornis* samples were collected from the Xisha Islands in 2018, inclusive of three samples per species per timepoint (including the non-acidified control group). All samples were found in tropical shallow reefs where the daily mean temperature was 23.2–29.2 °C. Before the experiments, all coral samples were kept whole and housed in our laboratory coral tank for at least three months, where the conditions were set to mimic their habitat in the South China Sea.

### Coral culture system

The coral samples were temporarily cultured in a standard RedSea® tank (redsea575, Red Sea Aquatics Ltd., London, UK) before and after micro-CT, following the Berlin method. The temperature was kept at 25 °C and the salinity was 1.025. The culture system was maintained using a Protein Skimmer (regal250s, Honya Co. Ltd., Shenzhen, China), a water chiller (tk1000, TECO Ltd., Taiwan, China), three coral lamps (AI®, Red Sea Aquatics Ltd., London, UK), two wave devices (VorTechTM MP40, EcoTech Marine Ltd., Bethlehem, USA), and a calcium reactor (Calreact 200, Honya Co. Ltd., Shenzhen, China). This coral culture system is the husbandry setup for the full duration of the experiment. In the experiment, we utilized four sets of such cultivation systems, with three sets used for parallel experimental groups to simulate ocean acidification, and the remaining set used for the non-acidified control group. The corals were fragmented into uniform-sized pieces for the experiment.

### Simulation of ocean acidification

Carbonate chemistry in the coral tank was manipulated by bubbling CO_2_ through the water to reduce the control pH (pH 8.0–8.2) to the target value of pH 7.8, and the changing process of carbonate chemistry in coral tank can be found in Table S5. The experiment utilized ion concentration meters and pH meters for parameter measurements (pH, Ca^2+^, CO_3_^2−^, HCO ^−^). Meanwhile, we used a microfluidic device to periodically release quantities of acetic acid into the coral tank to make the pH fluctuate regularly between 7.8 during the day and 7.6 during the night. The pH of the tank was monitored continuously by electrodes linked to a monitoring system (Figure S28).

### Micro-CT test

Coral samples from the South China Sea were analyzed using 3D models constructed with a 230 kV latest-generation X-ray microfocus computed tomography system (Phoenix v|tome|x m; General Electric, GE; at Yinghua NDT, Shanghai, China; Table S4). Two-dimensional image reconstructions of each specimen from the matrices of scan slices were assembled using proprietary software from GE. We conducted micro-CT tests on three samples per species at each timepoint, with the samples being collected from different colonies and exhibiting nearly identical sizes.

### Micro-CT reconstructions

Slice data derived from the scans were then analyzed and manipulated using 3D reconstruction software. Polyp-canal system reconstructions and skeleton to void ratio measurements were performed using VG Studio Max^15,26^ (v3.3.0). The 3D reconstructions were created following the method previously described. Images of the reconstructions were exported from Mimics and VG Studio Max and finalized in Adobe Photoshop CC 2019 and Adobe Illustrator CC 2019.

### Skeleton to void ratio measurement

The calculation of the skeletal matter to void volumetric ratio of coral samples, referred to as “skeleton to void ratio”, was conducted using VG Studio Max 3.3^27^. Initially, the “surface determination” function was utilized to differentiate between the areas of reconstructed skeleton and canal system in the colony. Subsequently, the volume of the reconstructed skeleton was calculated and the “erode/dilate” mode was employed to encompass the entire area of both the skeleton and canals. Following this, the “porosity/inclusion analysis module” was used to reconstruct the lumen of the canal system for volume calculation. Consequently, with volumes obtained for both the skeleton and internal canal system, we were able to determine a skeleton-void ratio for each sample. Three samples per species per timepoint were measured along with three technical replicates per sample in this study.

### SEM-EDS test

For the SEM-EDS analysis, 3 mm × 3 mm × 1 mm cubes were extracted from the central region of each coral colony utilized in the micro-CT examination. To ensure the precision of the SEM analysis, conductive coatings were meticulously applied to polished 3 mm × 3 mm cross-sections of each sample. Subsequently, the coral samples underwent SEM imaging to capture cross-sectional views of their skeletons^28^. Following this, three random rectangular areas measuring 0.15 mm × 0.10 mm were selected from the SEM images of each sample for EDS scanning^29^. The atomic ratios of Ca, O and C within each area were determined and averaged across all samples for statistical analysis purposes. A total of three samples per species per timepoint were analyzed using SEM-EDS.

### RNA-seq test

#### 1) Sampling

The simulation device was turned on, then when the tank pH stabilized at 7.6–7.8, the test started at Day 1. Before the simulation, samples were denoted as “Day 0”, then denoted according to the experimental day.

#### 2) Total RNA extraction

Three samples per species per timepoint were measured along with three technical replicates per sample in this test, and the three samples from each colony were treated independently for the transcriptome sequencing. In each coral, triplicate biological samples were isolated from three healthy branches in the same colony to ensure that enough high-quality RNA (> 15 µg) could be obtained for a PacBio cDNA library and three Illumina cDNA libraries. All the RNA extraction procedures followed the manufacturer’s instructions. The total RNA was isolated with TRIzol^®^ LS Reagent (Thermo Fisher Scientific, 10296028, Waltham, MA, USA) and treated with DNase I (Thermo Fisher Scientific, 18068015, Waltham, MA, USA). The high-quality mRNA was isolated with a FastTrack MAG Maxi mRNA Isolation Kit (Thermo Fisher Scientific, K1580-02, Waltham, MA, USA). The RNA extraction procedure was performed according to the following instructions: (1) grind the coral samples into small pieces (submerged in liquid nitrogen at all times); (2) add TRIzol^®^ LS reagent at a sample-to-reagent ratio of about 1:3; (3) let the samples stand and thaw naturally; (4) continue adding TRIzol^®^ LS reagent until the samples are dissolved, and dispense into 50 mL centrifuge tubes; (5) centrifuge at 4 and 3000 rpm for 5–15 min; (6) dispense the supernatant into 50 mL centrifuge tubes; (7) add BCP (Molecular Research Center, BP 151, Cincinnati, OH, USA) to the centrifuge tubes at a sample-to-reagent ratio of about 5:1, shake well and stand for 10 min; (8) centrifuge at 4 and 10,500 rpm for 15 min; (9) remove the supernatant, add an equal volume of isopropanol (Amresco, 0918-500ML, Radnor, PA, USA) and mix well, stand overnight at –20; (10) centrifuge at 4 and 10,500 rpm for 30 min, discard the supernatant; (11) rinsed twice with 75% ice ethyl alcohol, pure (Sigma-Aldrich, E7023-500ML, Taufkirchen, München, Germany). Finally, extract three samples of each coral in equal amounts (total > 10 µg) and mix for PacBio full-length transcriptome sequencing. The remainder (> 1.5 µg per sample) was used for Illumina sequencing.

#### 3) Total RNA quality testing

Before establishing the library, the quality of total RNA must be tested. RNA degradation and contamination were monitored by 1% agarose gels electrophoresis; RNA purity (OD260/280 ratio) was checked using a NanoPhotometer^®^ spectrophotometer (IMPLEN, CA, USA); RNA concentration was quantified using a Qubit® RNA Assay Kit in Qubit^®^ 2.0 Fluorometer (Life Technologies, CA, USA); and RNA integrity was assessed using the RNA Nano 6000 Assay Kit of the Agilent Bioanalyzer 2100 system (Agilent Technologies, CA, USA).

#### 4) Illumina cDNA library construction and sequencing

A total amount of 1.5 µg RNA per sample was used as input material for the RNA sample preparations. Sequencing libraries were generated using NEBNext^®^ Ultra™ RNA Library Prep Kits (E7530L) for Illumina^®^ (NEB, Ipswich, MA, USA) following the manufacturer’s recommendations and index codes were added to attribute sequences to each sample. Briefly, mRNA was purified from total RNA using poly-T oligo-attached magnetic beads. Fragmentation was carried out using divalent cations under elevated temperature in NEBNext First Strand Synthesis Reaction Buffer (5×). First-strand cDNA was synthesized using random hexamer primer and M-MuLV Reverse Transcriptase (RNase H^−^). Second-strand cDNA synthesis was subsequently performed using DNA Polymerase I and RNase H. Remaining overhangs were converted into blunt ends via exonuclease/polymerase activities. After adenylation of 3’ ends of DNA fragments, the NEBNext Adaptor with a hairpin loop structure was ligated to prepare for hybridization. To select cDNA fragments preferentially of 250–300 bp in length, the library fragments were purified with the AMPure XP system (Beckman Coulter, Beverly, USA). Then, 3 µL USER Enzyme (NEB, USA) was used with size-selected, adaptor-ligated cDNA at 37 for 15 min followed by 5 min at 95 before PCR. Then, PCR was performed with Phusion High-Fidelity DNA polymerase, Universal PCR primers and Index (X) Primer. Finally, the PCR products were purified (AMPure XP system) and library quality was assessed on the Agilent Bioanalyzer 2100 system. Clustering of index-coded samples was performed on a cBot Cluster Generation System using a TruSeq PE Cluster Kit v3-cBot-HS (Illumia) according to the manufacturer’s instructions. After cluster generation, the library preparations were sequenced on an Illumina HiSeq X Ten platform and paired-end reads were generated.

#### 5) PacBio cDNA library construction and sequencing

An isoform sequencing (Iso-Seq) library was prepared according to the Iso-Seq protocol using the Clontech SMARTer^®^ PCR cDNA Synthesis Kit (Clontech Laboratories, now Takara Laboratories, 634926, Mountain View, CA, USA) and the BluePippin Size Selection System protocol as described by Pacific Biosciences (PN 100-092-800-03). Briefly, Oligo(dT)-enriched mRNA was reversely transcribed to cDNA by a SMARTer PCR cDNA Synthesis Kit. The synthesized cDNA was then amplified by polymerase chain reaction (PCR) using the BluePippin Size Selection System protocol. The Iso-Seq library was constructed by full-length cDNA damage repair, terminal repair and attaching SMRT dumbbell adapters. The sequences of the unattached adapters at both ends of the cDNA were removed by exonuclease digestion. The obtained cDNA was combined with primers and DNA polymerase to form a complete SMRT bell library. While the library was qualified, the PacBio Sequel II platform was used for sequencing based on the effective concentration and data output requirements of the library.

#### 6) Data Filtering and Processing

The Illumina sequencing raw reads in fastq format were first processed using in-house Perl scripts. In this step, clean data were obtained by removing reads containing adapter or ploy-N, and low-quality reads from raw data. At the same time, the Q20, Q30, GC-content and sequence duplication levels of the clean data were calculated. All the downstream analyses were based on clean data with high quality.

The PacBio sequencing raw data were processed by SMRTlink v8.0 software. A circular consensus sequence (CCS) was generated from subread BAM files using the parameters: min_length 50, min_passes 1, max_length 15,000. CCS.BAM files were output, which were then classified into full-length and non-full-length reads using lima, removing polyA using refine. Full-length fasta files were produced and then fed into the cluster step, which performed isoform-level hierarchical clustering (*n*log(*n*)), followed by final Arrow polishing using the settings hq_quiver_min_accuracy 0.99, bin_by_primer false, bin_size_kb 1, qv_trim_5p 100, qv_trim_3p 30.

#### 7) Coral and Symbiodiniaceae sequences separation

Aligned consensus reads to coral or Symbiodiniaceae reference genomes were performed using GMAP v2017-06-20 software^30^. The sequences mapped to Symbiodiniaceae reference genomes belonged to Symbiodiniaceae sequences, while sequences mapped to coral reference genomes belonged to coral sequences.

#### 8) Correction and de-redundancy

The RNA-seq data sequenced by the Illumina HiSeq X Ten platform was used to correct additional nucleotide errors in polish consensus sequences obtained in the previous step with LoRDEC v0.7 software^31^. Using CD-HIT v4.6.8 software (parameters: -c 0.95 -T 6 -G 0 - aL 0.00 -aS 0.99), all redundancies were removed in corrected consensus reads to acquire final full-length transcripts and unigenes for subsequent bioinformatics analysis^32^.

#### 9) Gene functional annotation

Gene functions were annotated using the following databases: NT (NCBI non-redundant nucleotide sequences); NR (NCBI non-redundant protein sequences); Pfam (Protein family); KOG/COG (Clusters of Orthologous Groups of proteins); Swiss-Prot (A manually annotated and reviewed protein sequence database); KEGG (Kyoto Encyclopedia of Genes and Genomes); and GO (Gene Ontology). We used BLAST 2.7.1+ software^33^ with the e-value ‘1e-5’ for NT database analysis, Diamond v0.8.36 BLASTX software^34^ with the e-value ‘1e-5’ for NR, KOG, Swiss-Prot and KEGG databases analyses, and the HMMER 3.1 package^35^ for Pfam database analysis.

#### 10) Gene structure analysis

ANGEL v2.4 software^36^ was used to predict protein CDSs (coding sequences). We used the same species or closely related species-confident protein sequences for ANGEL training and then ran the ANGEL prediction for the given sequences. Usually, the TFs were identified based on the Pfam files of TF families in the AnimalTFDB 3.0 database^37^; however, corals were not included in this database, so we identified coral TFs based on the Pfam files of TF families using the hmmsearch program in the HMMER 3.1 package. The SSR of the transcriptomes was identified using MISA v1^38^. We used four tools—CNCI v2^39^, CPC2 v0.1^40^, PfamScan v1.6^41^, and PLEK v1.2^42^—to predict the coding potential of the transcripts. Transcripts predicted with coding potential by either/all of the above three tools were filtered out, and those lacking coding potential were our candidate set of lncRNAs.

#### 11) Gene expression quantification

The full-length transcriptome obtained above was used as the reference background, and then the clean reads of each sample obtained by Illumina sequencing were mapped to it using bowtie2 v2.3.4 software^43^. The alignment results were estimated by RSEM v1.3.0 software^44^ to obtain the read count values for each transcript, which were then transferred to FPKM for analysis of gene expression levels. Pearson correlation coefficients were used to analyze the relationships among samples.

#### 12) Gene differential expression analysis

The unigenes with the same annotation results in the NR database were merged to form a new read count expression matrix. Differential expression analysis of two groups was performed using the DESeq2 R package (v1.30.1)^45^. DESeq2 provides statistical routines for determining differential expressions in digital gene expression data using a model based on the negative binomial distribution. The resulting *p*-values were adjusted using Benjamini and Hochberg’s approach for controlling the false discovery rate and were named *p*_adj_. Coral genes with *p*_adj_ < 0.001 and |log2(FoldChange)| ≥ 2 and 10 as determined by DESeq2 were assigned as differentially expressed. Venn diagrams were drawn using the VennDiagram R package (1.6.20) and GO classification bar charts were drawn using the ggplot2 R package (3.3.5)^46^.

## Supporting information

Manuscript

## Ethics

This study did not involve experiments on cephalopods or higher animals. All coral sample collection and processing were performed according to the local laws governing the welfare of invertebrate animals.

## Author Contributions

Y. L., C. H. and Z. L. conceived the project. X. L., and H. Z. collected the coral samples. Y. L. did the experiments, produced the figures, and wrote the paper. Y. L., Y. Z., J. C., C. H., and Z. L. edited the paper. All authors discussed and commented on the data.

## Data Availability Statement

Data produced in this study are available at the Sequence Read Archive (SRA) [https://www.ncbi.nlm.nih.gov/sra/] under accession numbers: SAMN16237127-SAMN16237130; SAMN16456055-SAMN16456058; SAMN16365802-SAMN16365813; SAMN16237439-SAMN16237462. Details can be checked in the Supplementary File.

