## Supplementary material for "Skeleton-forming responses of reef-building corals under ocean acidification": Manuscript

**Part 1 | Supplementary Figures**

**Part 2 | Supplementary Tables**

**Part 3 | Data Availability Statement**

**Part 1 | Supplementary Figures**

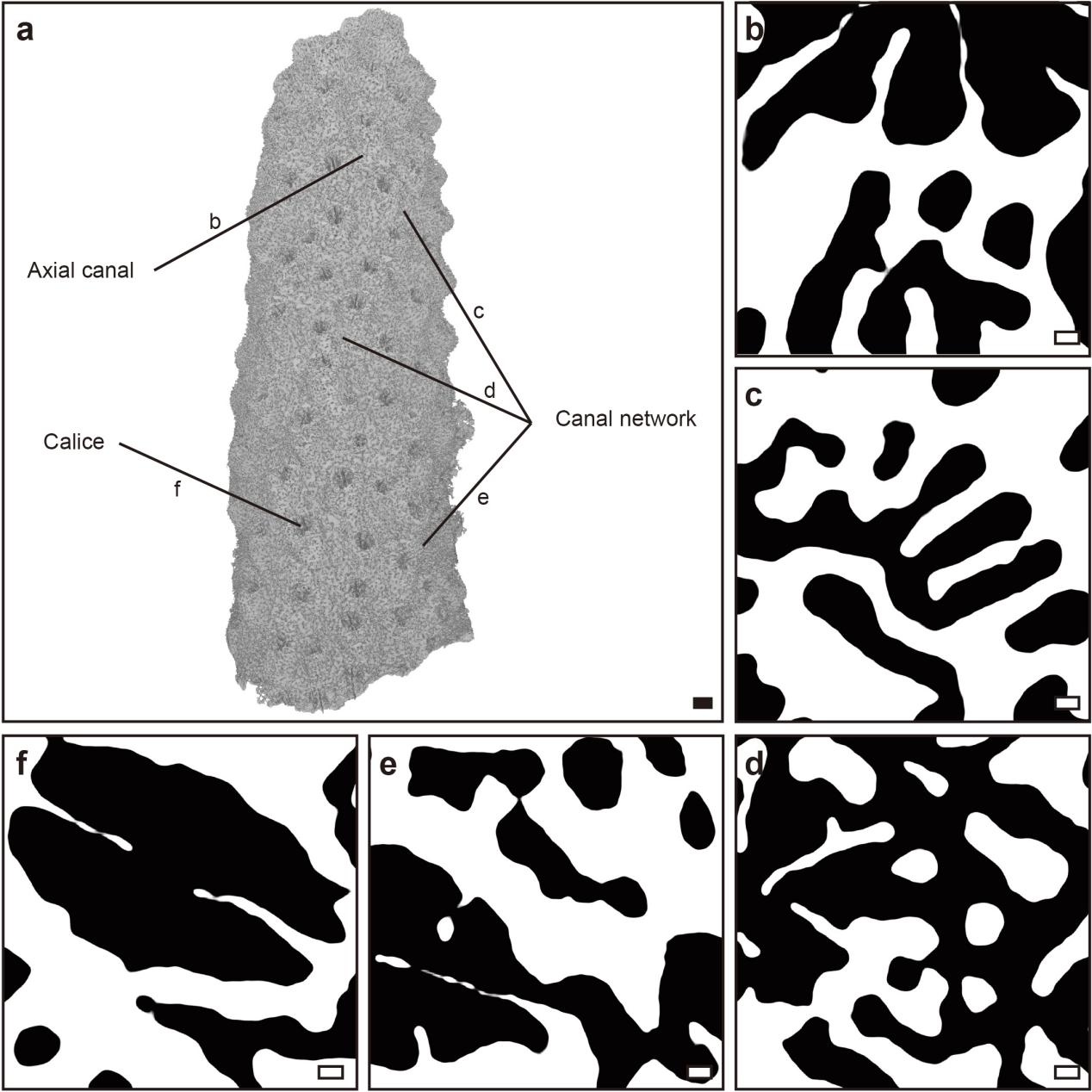

**Supplementary Figure 1 | Micro-CT reconstructions of *A. muricata* on Day 0.** Scale bars: a) 1 mm; b-f) 0.1 mm.

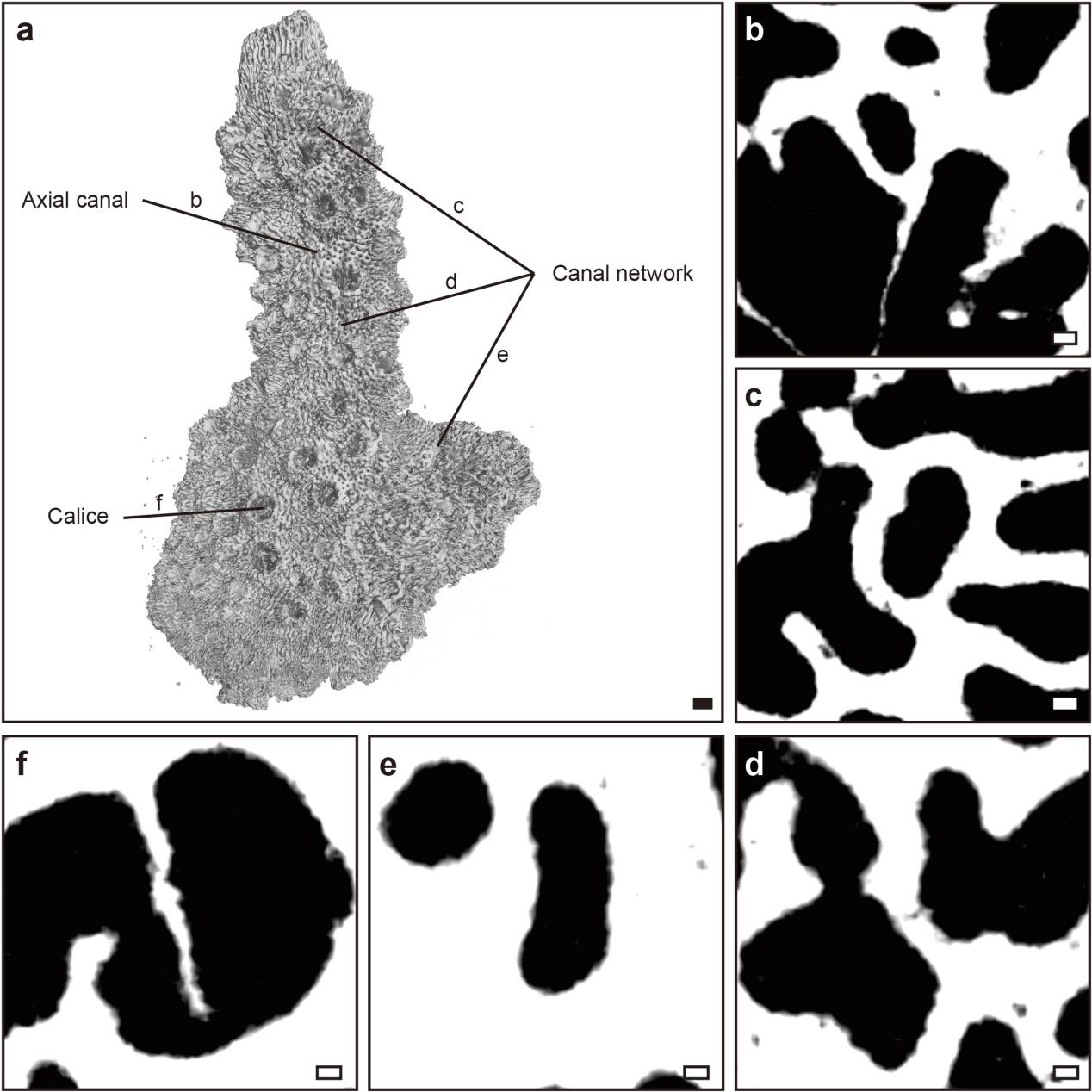

**Supplementary Figure 2 | Micro-CT reconstructions of *A. muricata* on Day 3.** Scale bars: a) 1 mm; b-f) 0.1 mm.

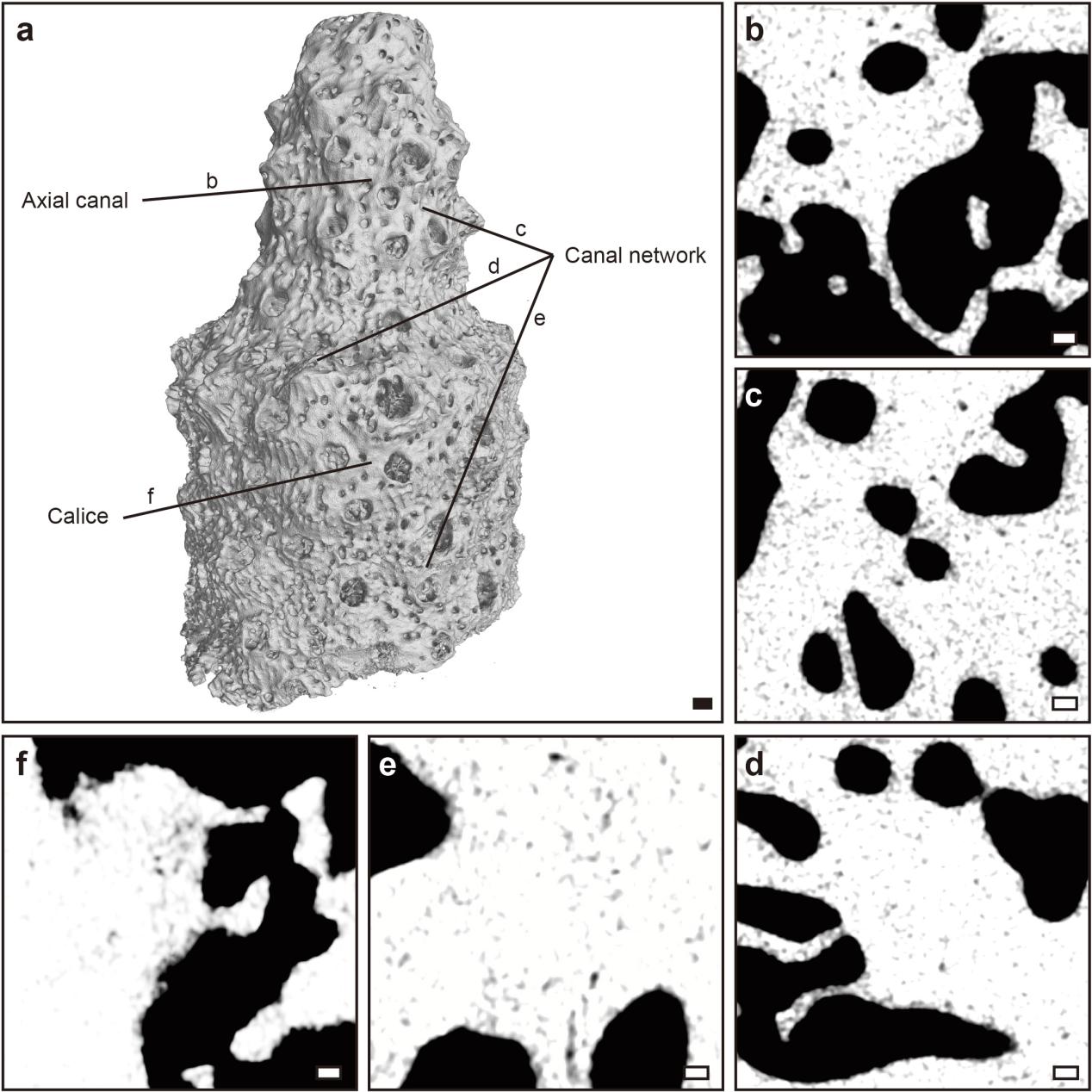

**Supplementary Figure 3 | Micro-CT reconstructions of *A. muricata* on Day 6.** Scale bars: a) 1 mm; b-f) 0.1 mm.

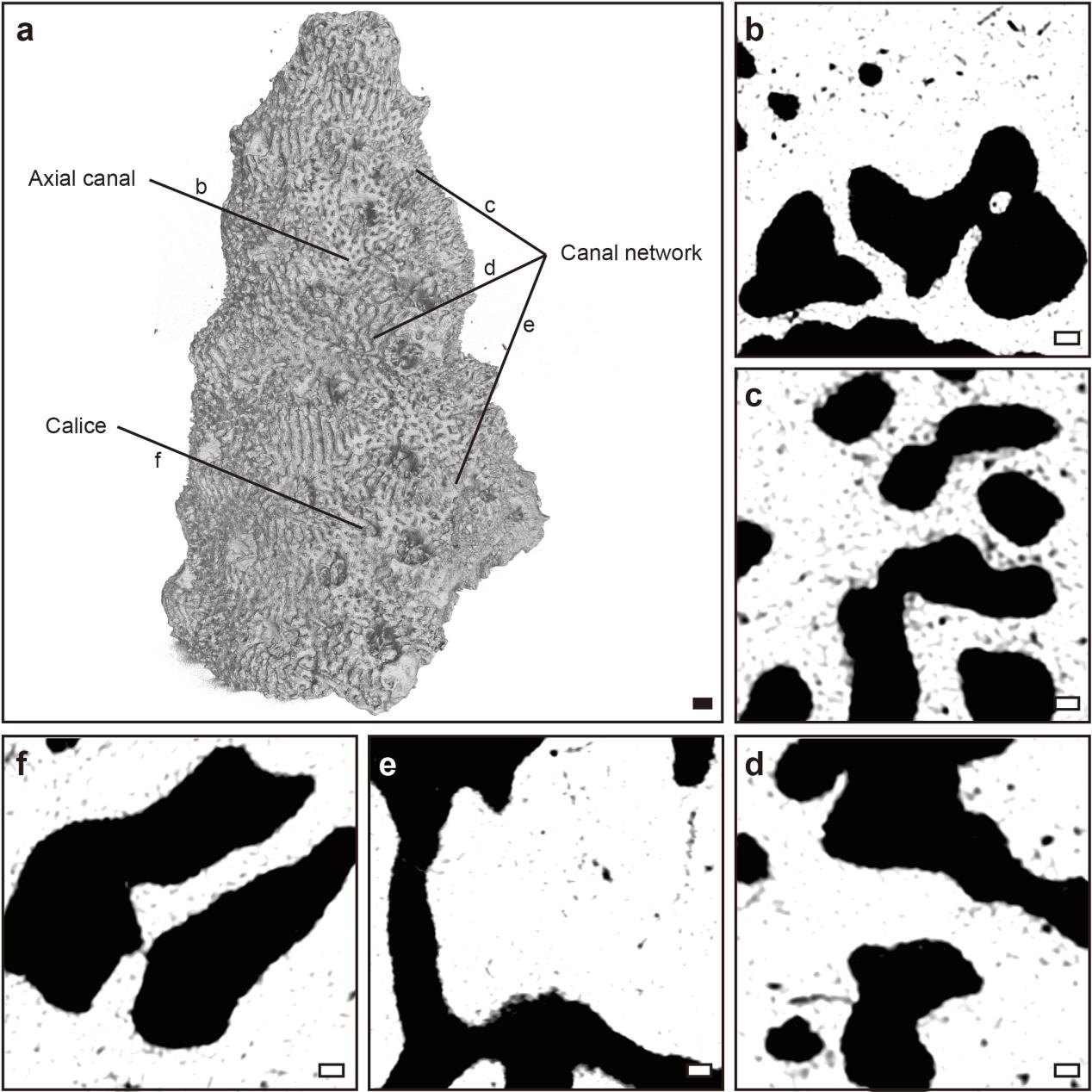

**Supplementary Figure 4 | Micro-CT reconstructions of *A. muricata* on Day 9.** Scale bars: a) 1 mm; b-f) 0.1 mm.

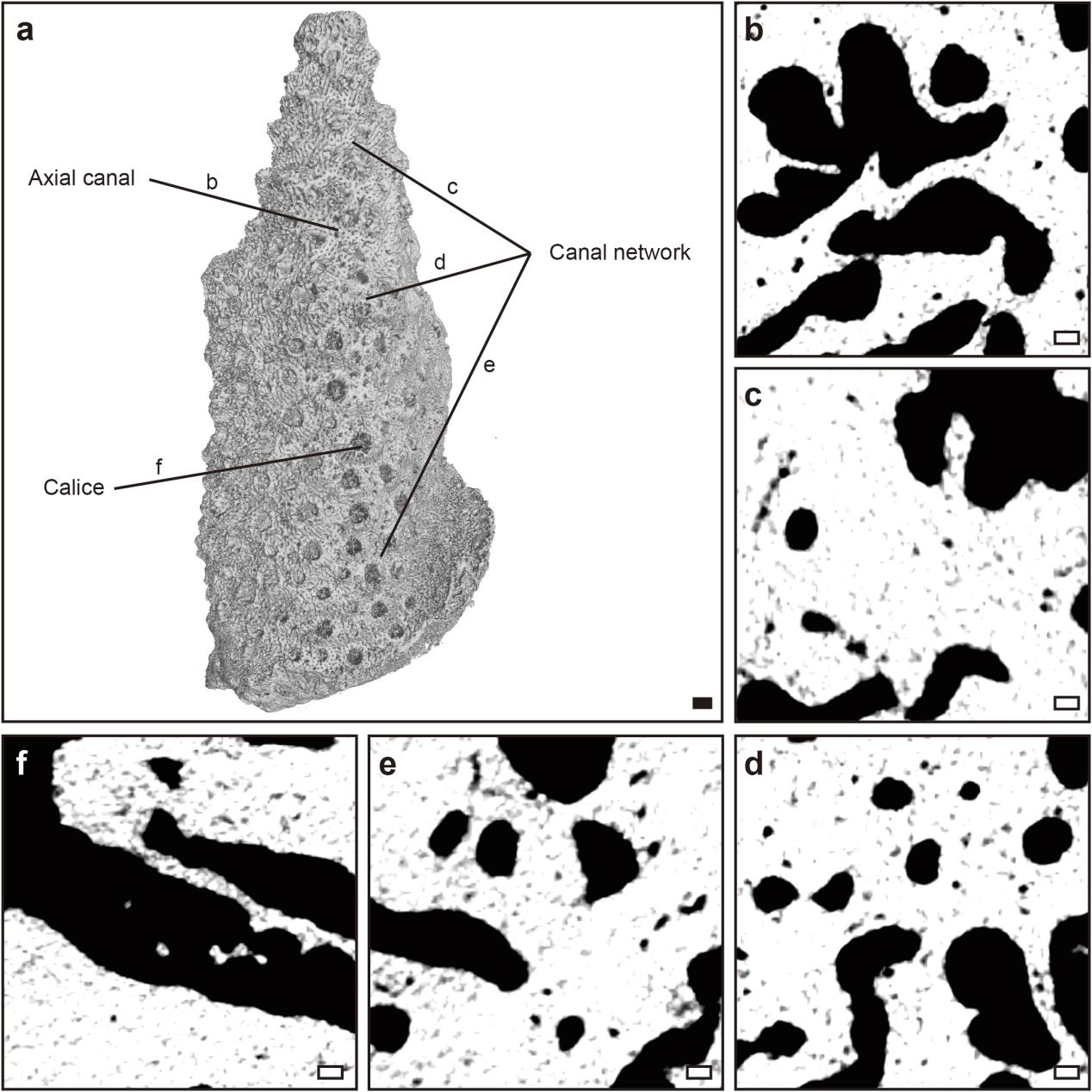

**Supplementary Figure 5 | Micro-CT reconstructions of *A. muricata* on Day 30.** Scale bars: a) 1 mm; b-f) 0.1 mm.

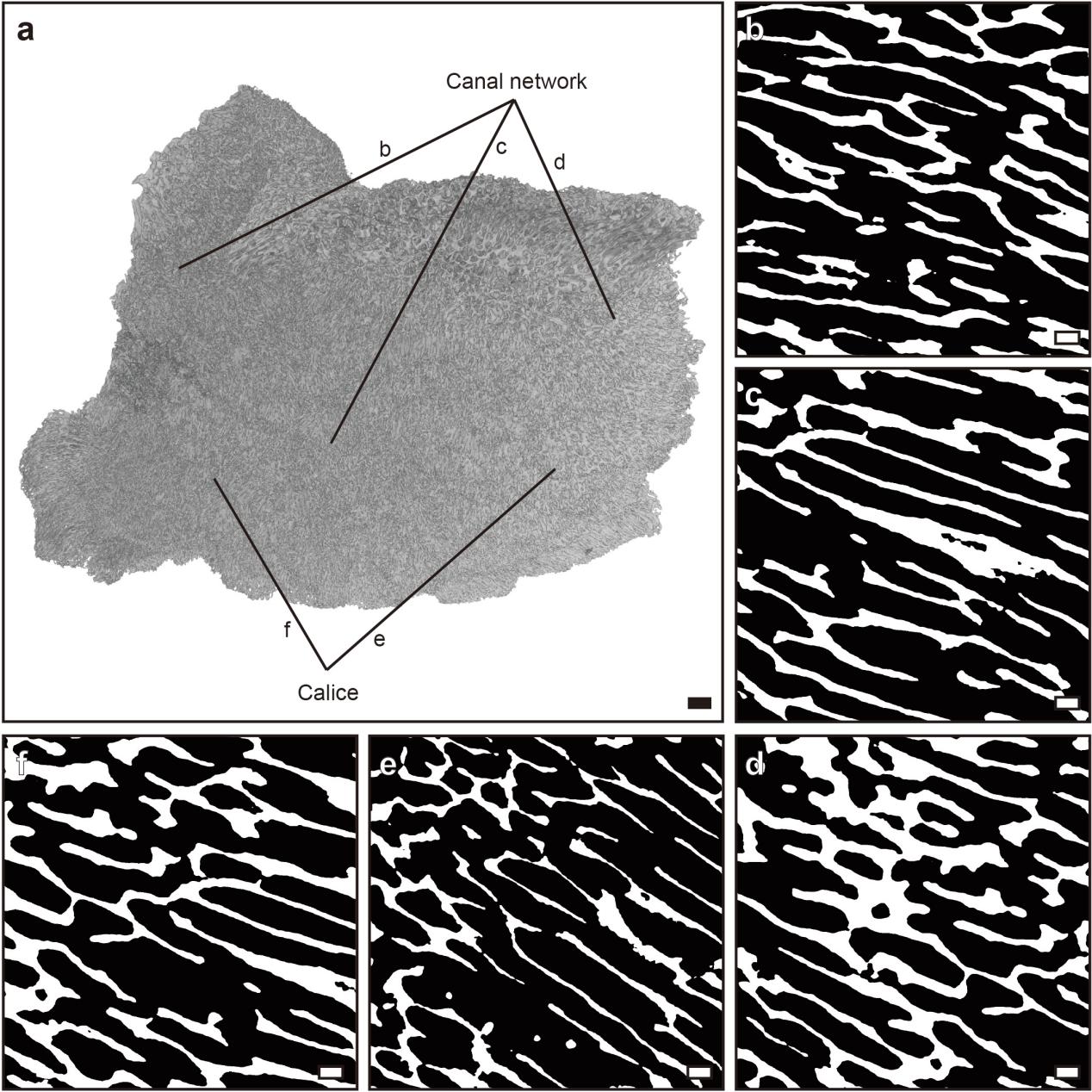

**Supplementary Figure 6 | Micro-CT reconstructions of *M. capricornis* on Day 0.** Scale bars: a) 1 mm; b-d) 0.1 mm; e,f) 0.2mm.

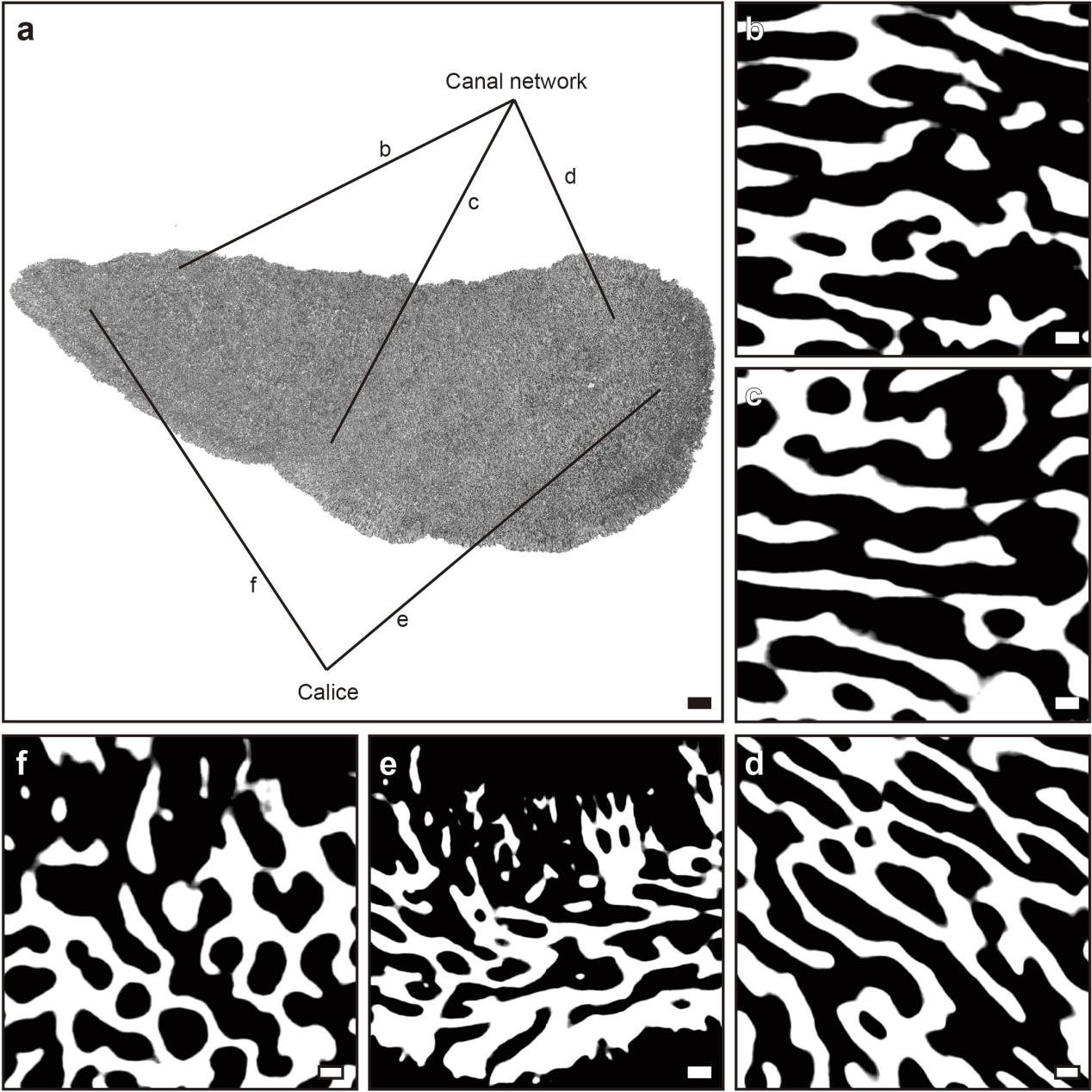

**Supplementary Figure 7 | Micro-CT reconstructions of *M. capricornis* on Day 3.** Scale bars: a) 1 mm; b-d) 0.1 μm; e) 0.2 mm; f) 0.1 mm.

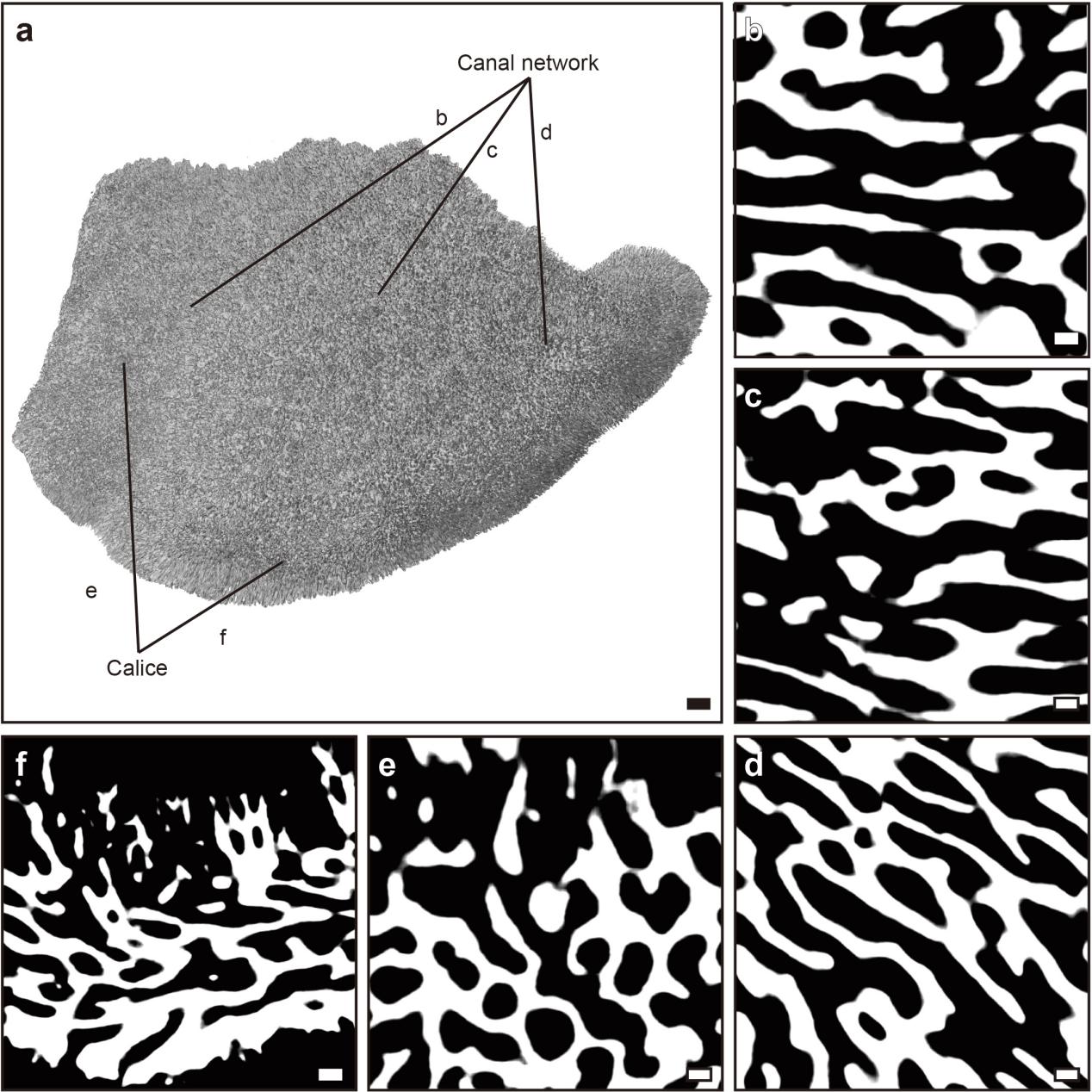

**Supplementary Figure 8 | Micro-CT reconstructions of *M. capricornis* on Day 6.** Scale bars: a) 1 mm; b-e) 0.1 mm; f) 0.2 mm.

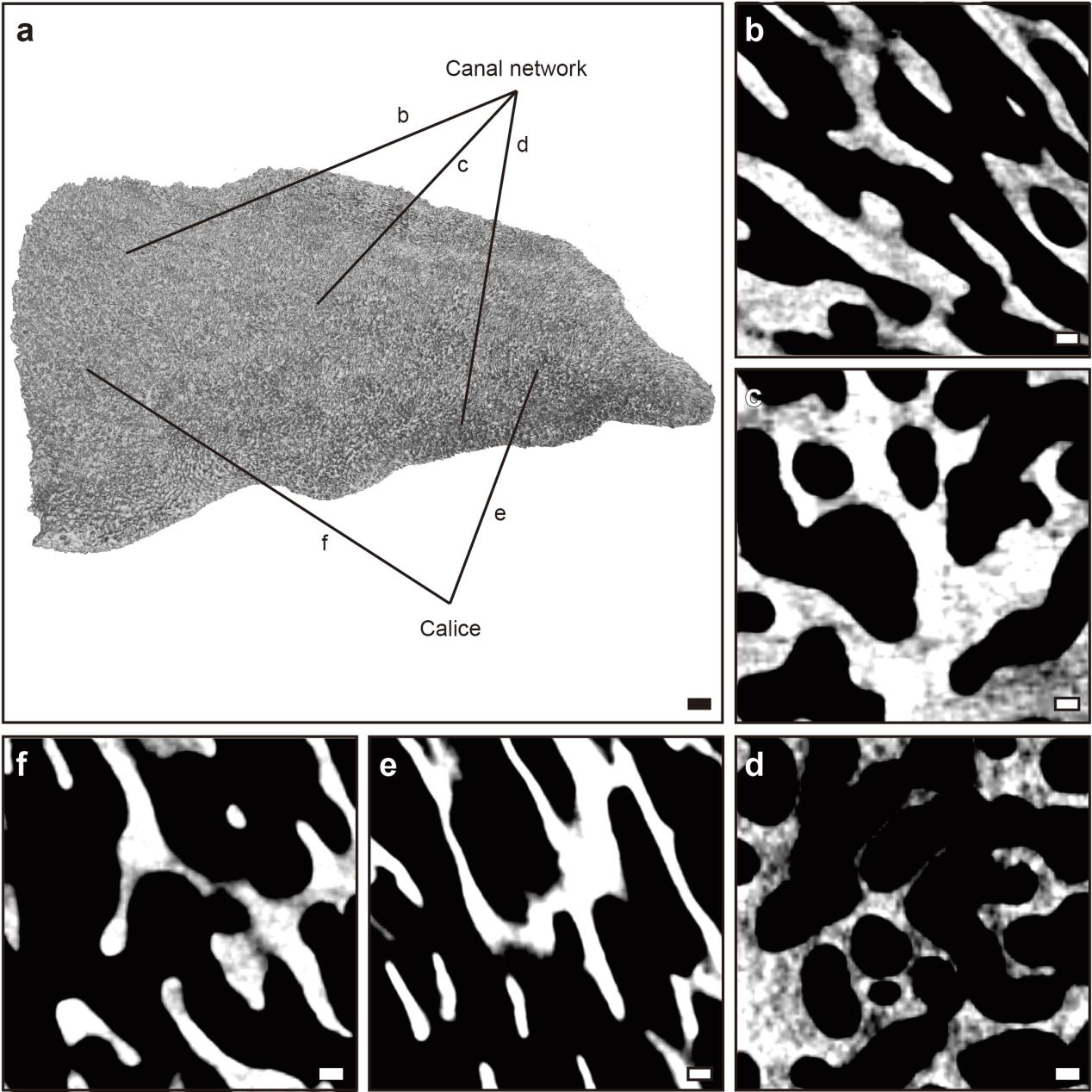

**Supplementary Figure 9 | Micro-CT reconstructions of *M. capricornis* on Day 9.** Scale bars: a) 1 mm; b-d) 0.1 mm; e,f) 0.2 mm.

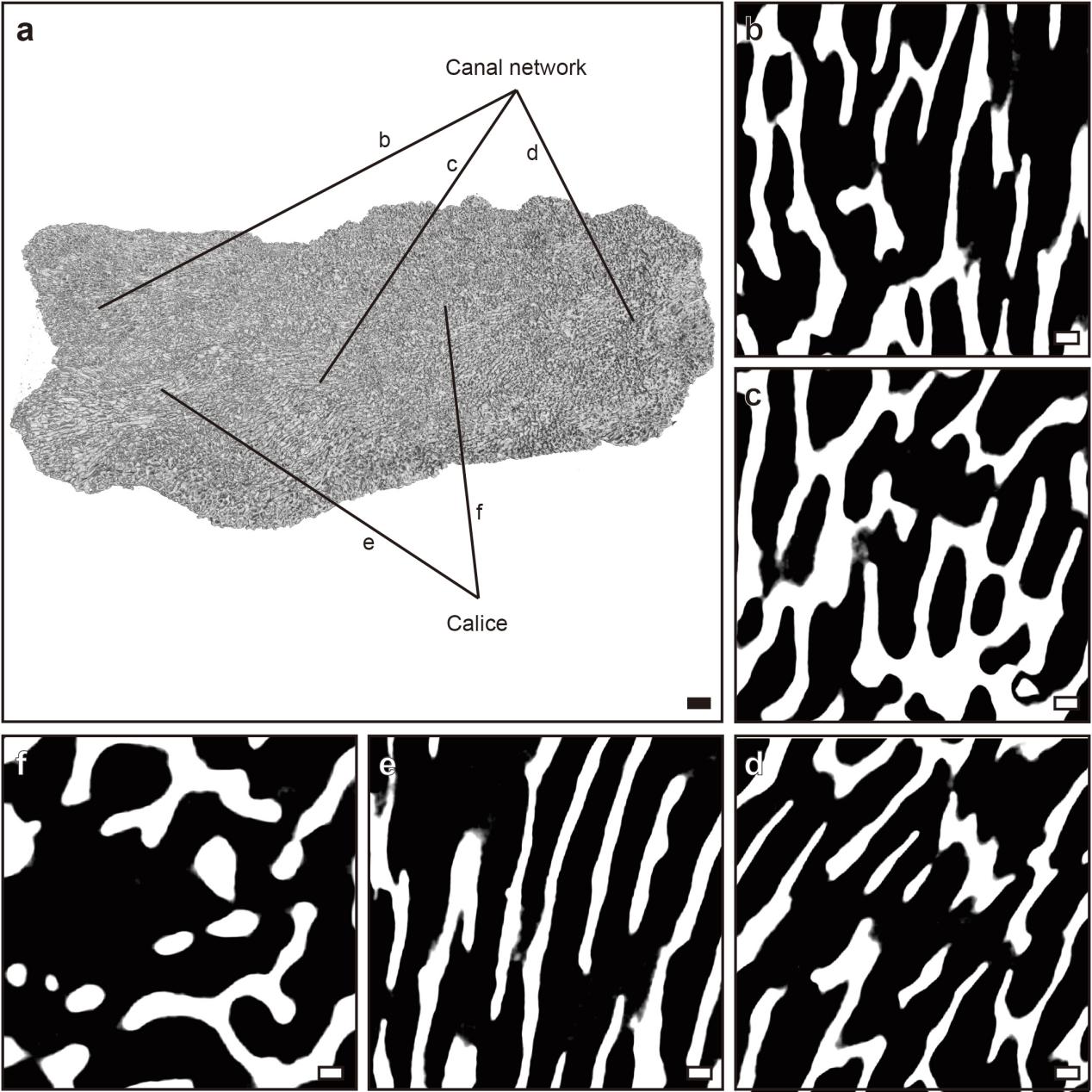

**Supplementary Figure 10 | Micro-CT reconstructions of *M. capricornis* on Day 30.** Scale bars: a) 1 mm; b-d) 0.1 mm; e) 0.2 mm; f) 0.1 mm.

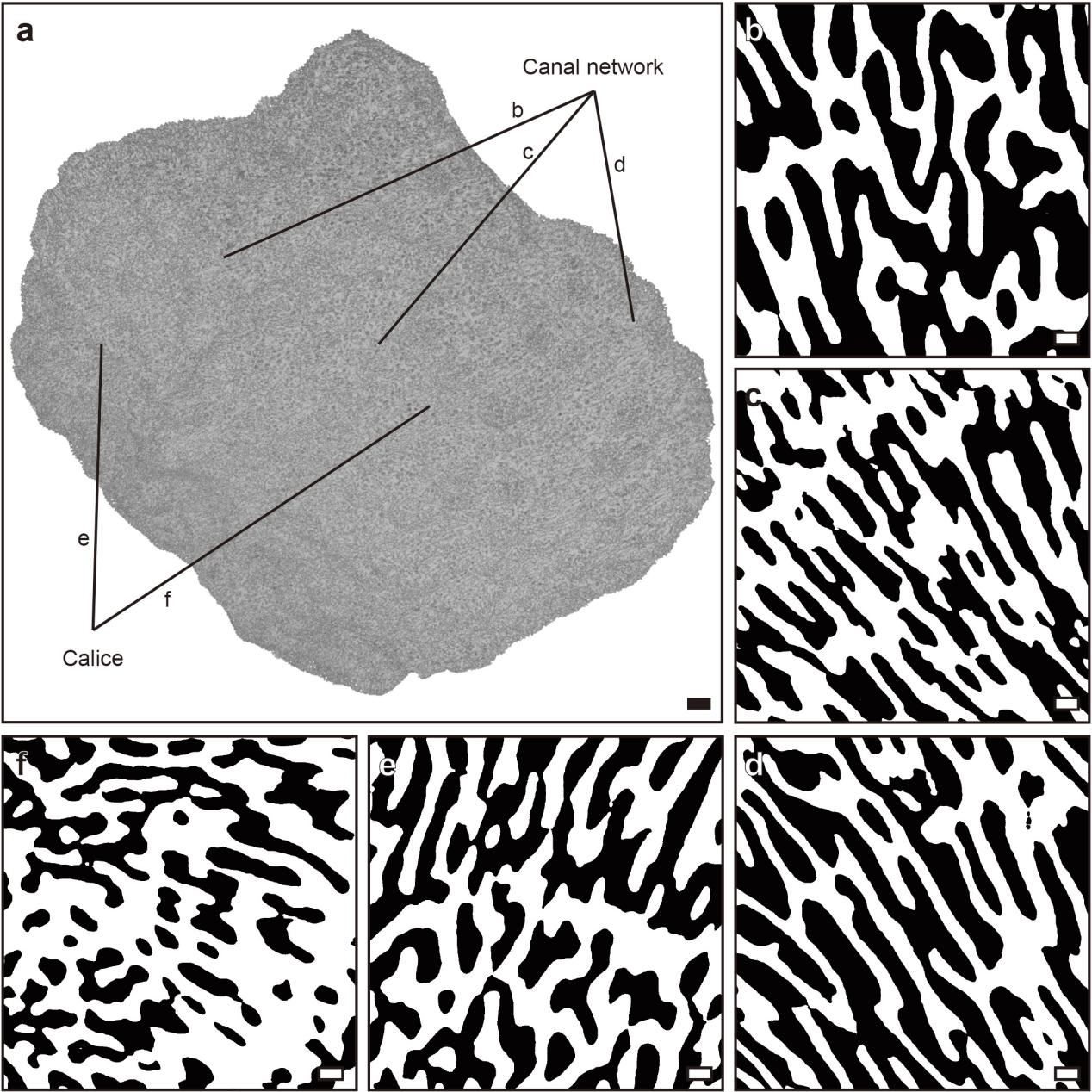

**Supplementary Figure 11 | Micro-CT reconstructions of *M. foliosa* on Day 0.** Scale bars: a) 1 mm; b-f) 0.1 mm.

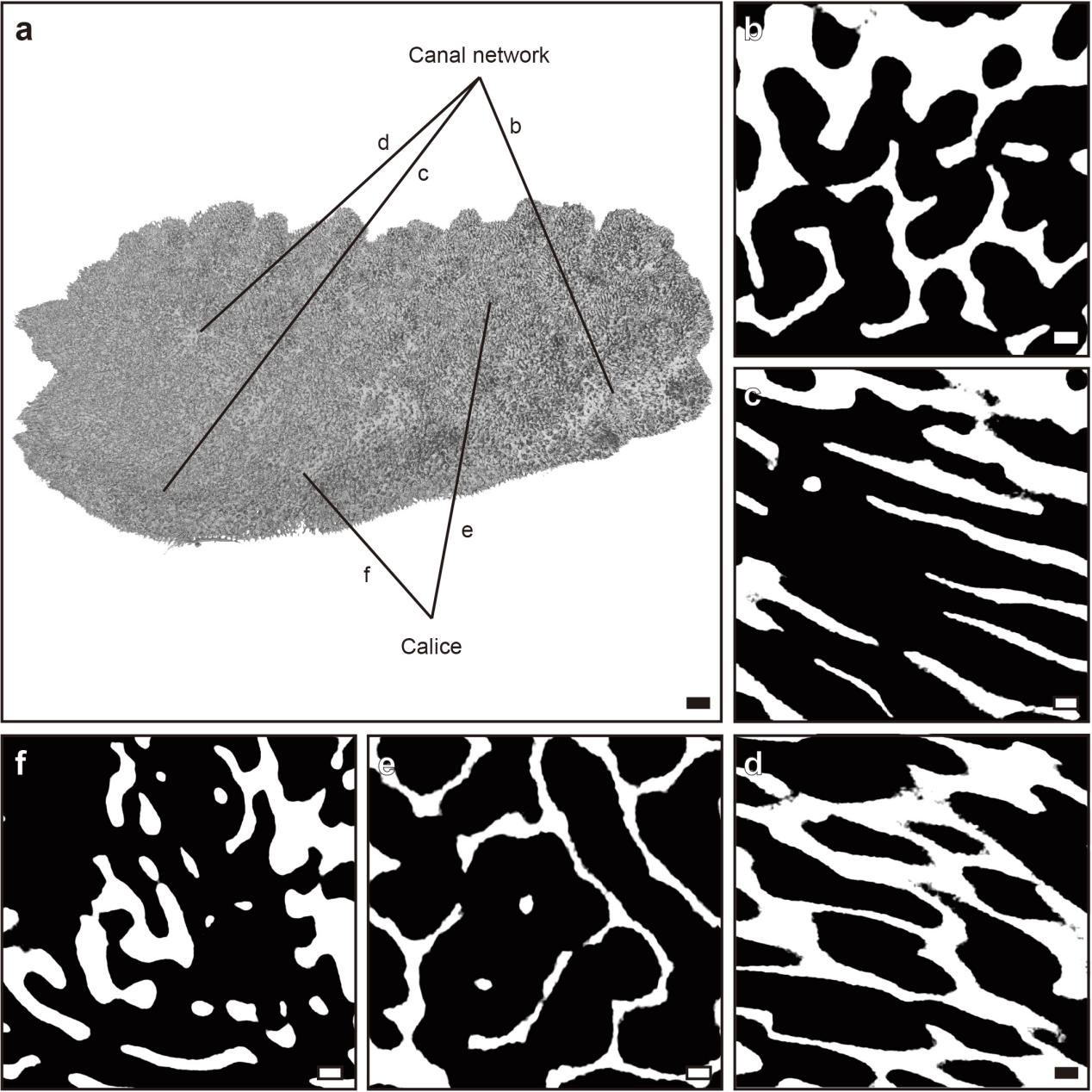

**Supplementary Figure 12 | Micro-CT reconstructions of *M. foliosa* on Day 3.** Scale bars: a) 1 mm; b-f) 0.1 mm.

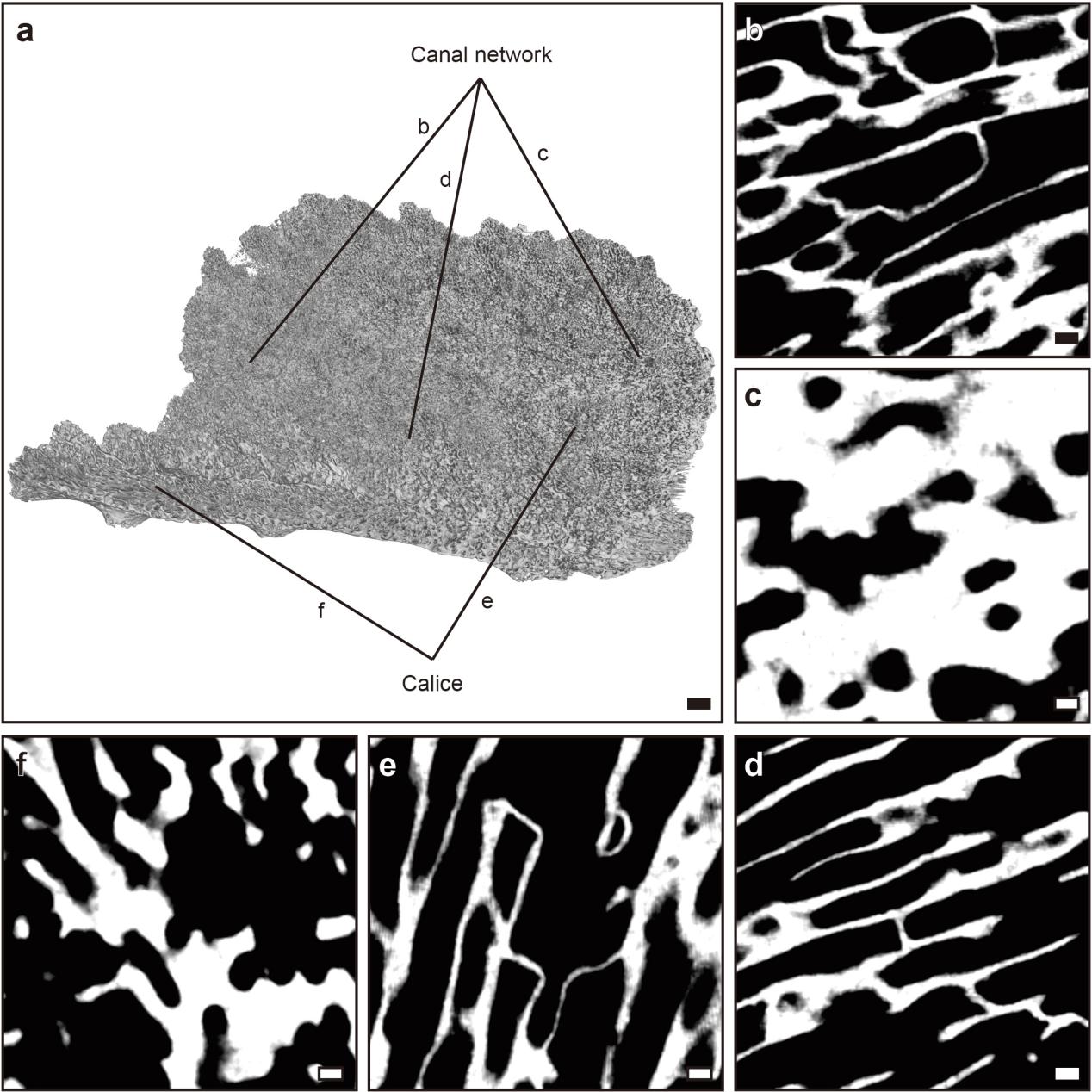

**Supplementary Figure 13 | Micro-CT reconstructions of *M. foliosa* on Day 6.** Scale bars: a) 1 mm; b-f) 0.1 mm.

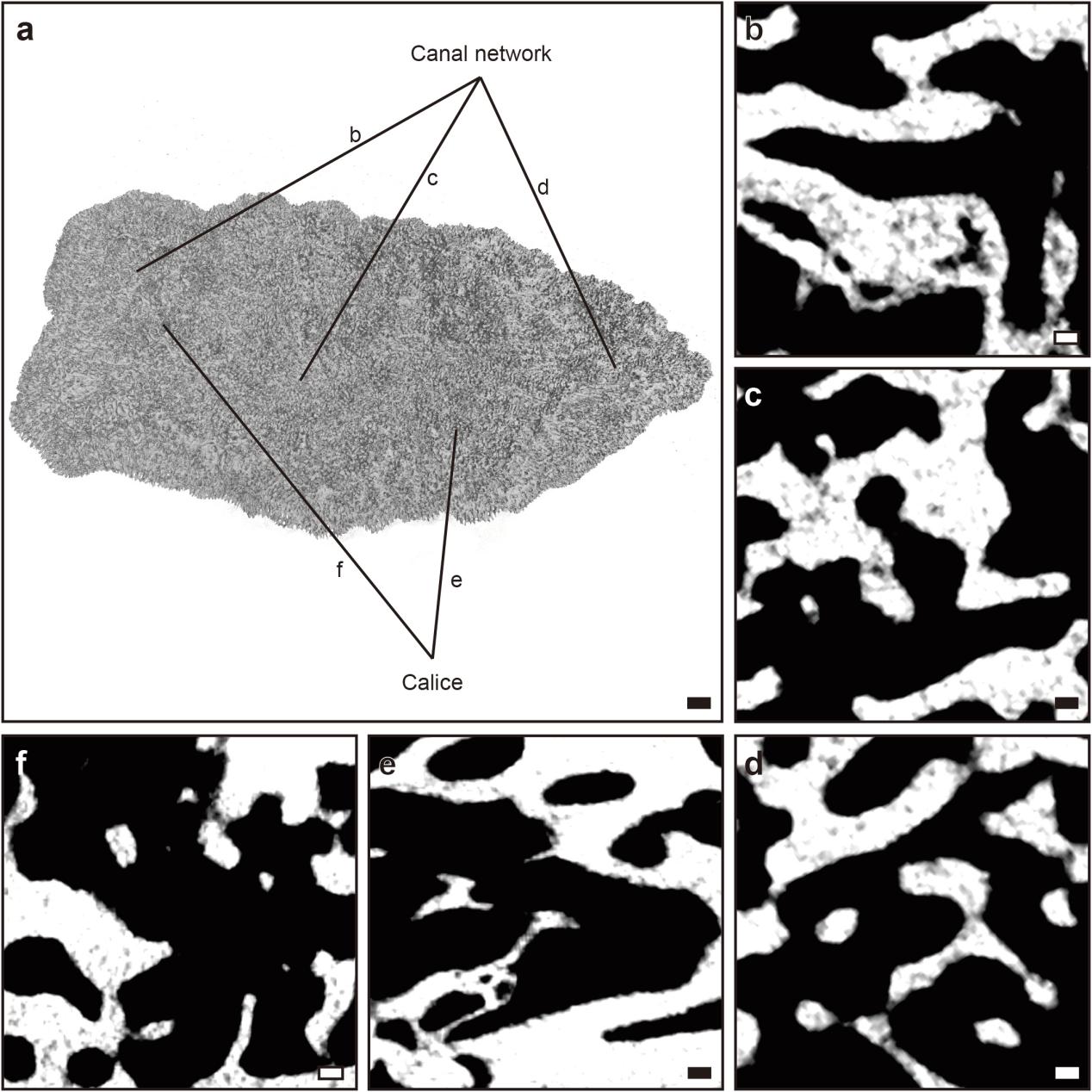

**Supplementary Figure 14 | Micro-CT reconstructions of *M. foliosa* on Day 9.** Scale bars: a) 1 mm; b-f) 0.1 mm.

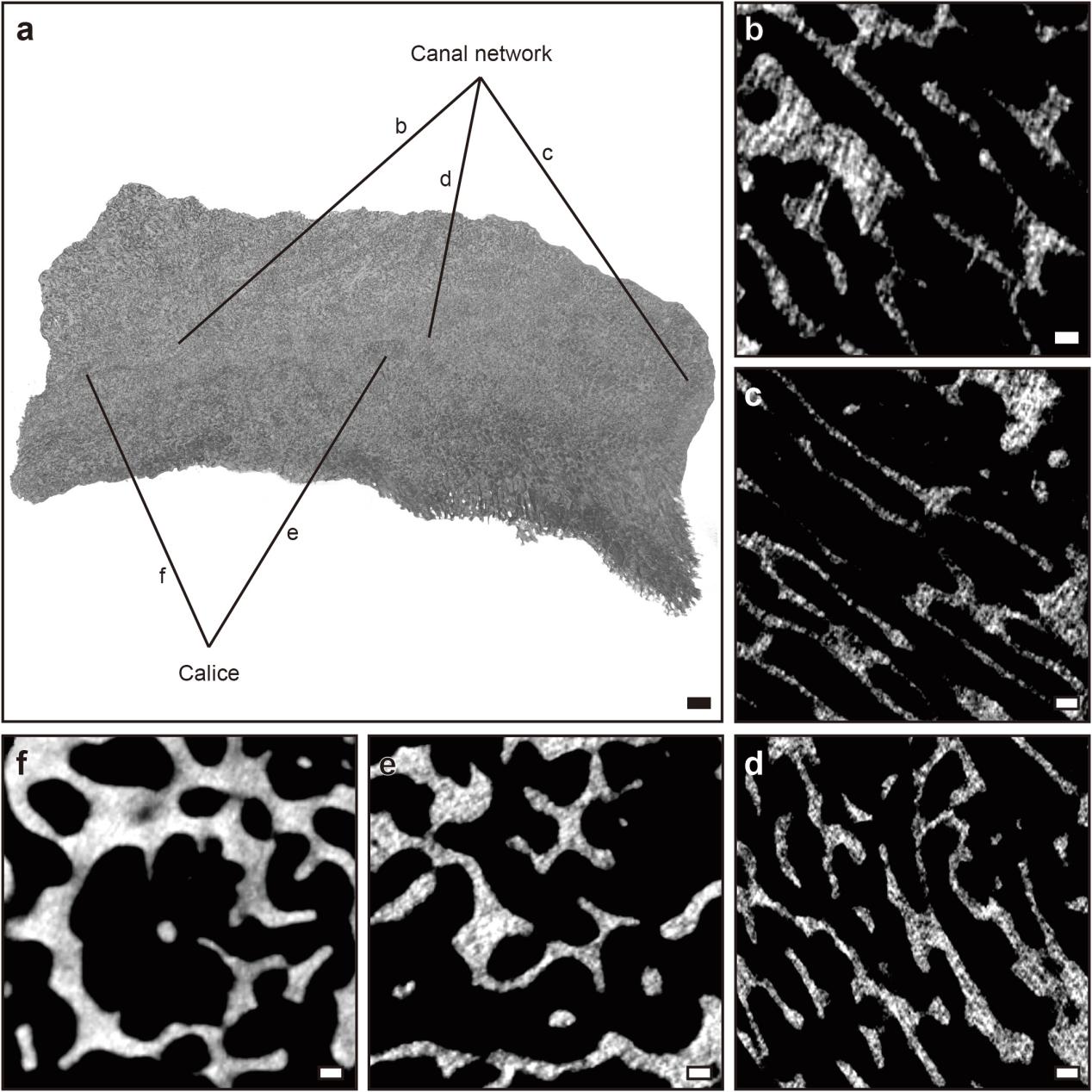

**Supplementary Figure 15 | Micro-CT reconstructions of *M. foliosa* on Day 30.** Scale bars: a) 1 mm; b-f) 0.1 mm.

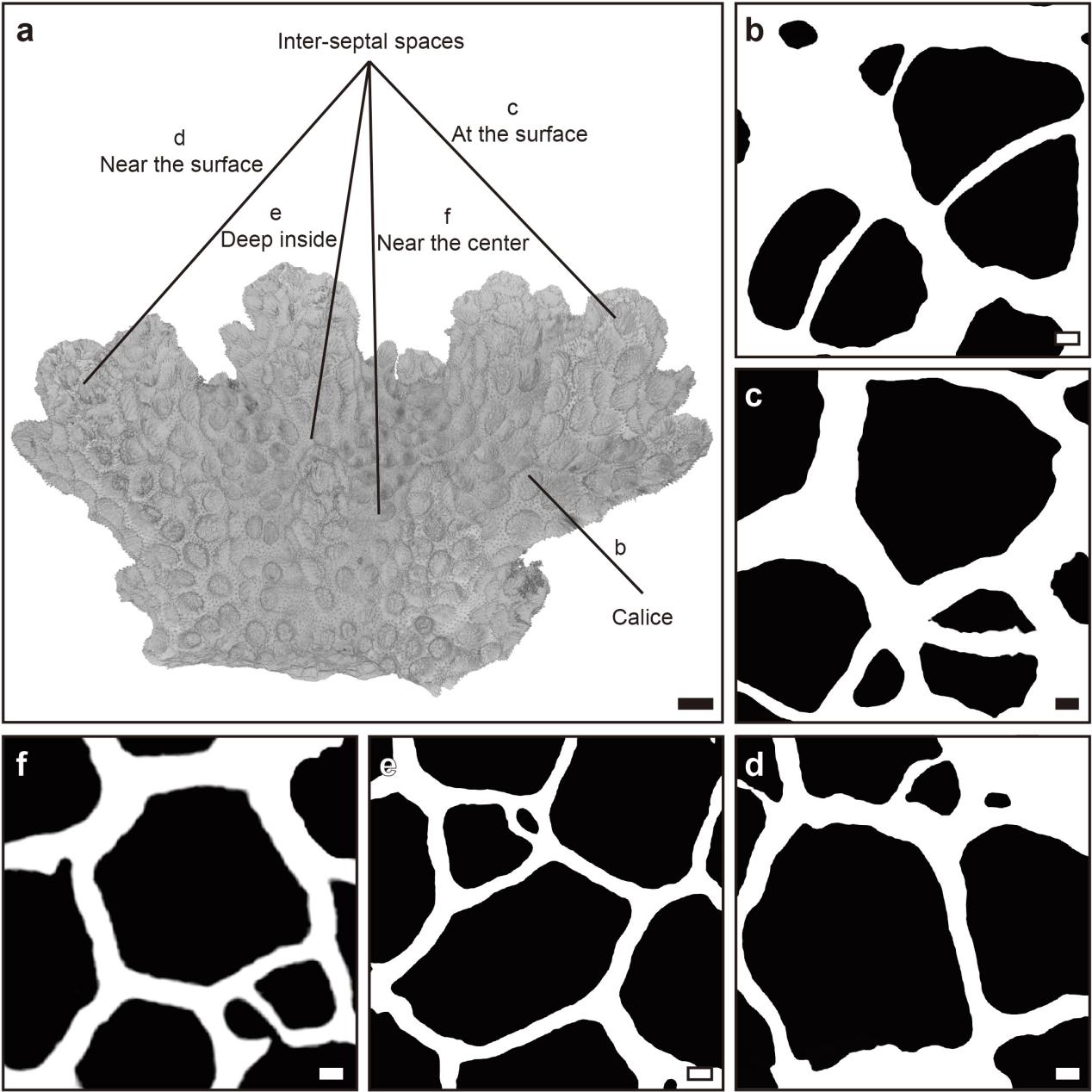

**Supplementary Figure 16 | Micro-CT reconstructions of *P. damicornis* on Day 0.** Scale bars: a) 1 mm; b-f) 0.1 mm.

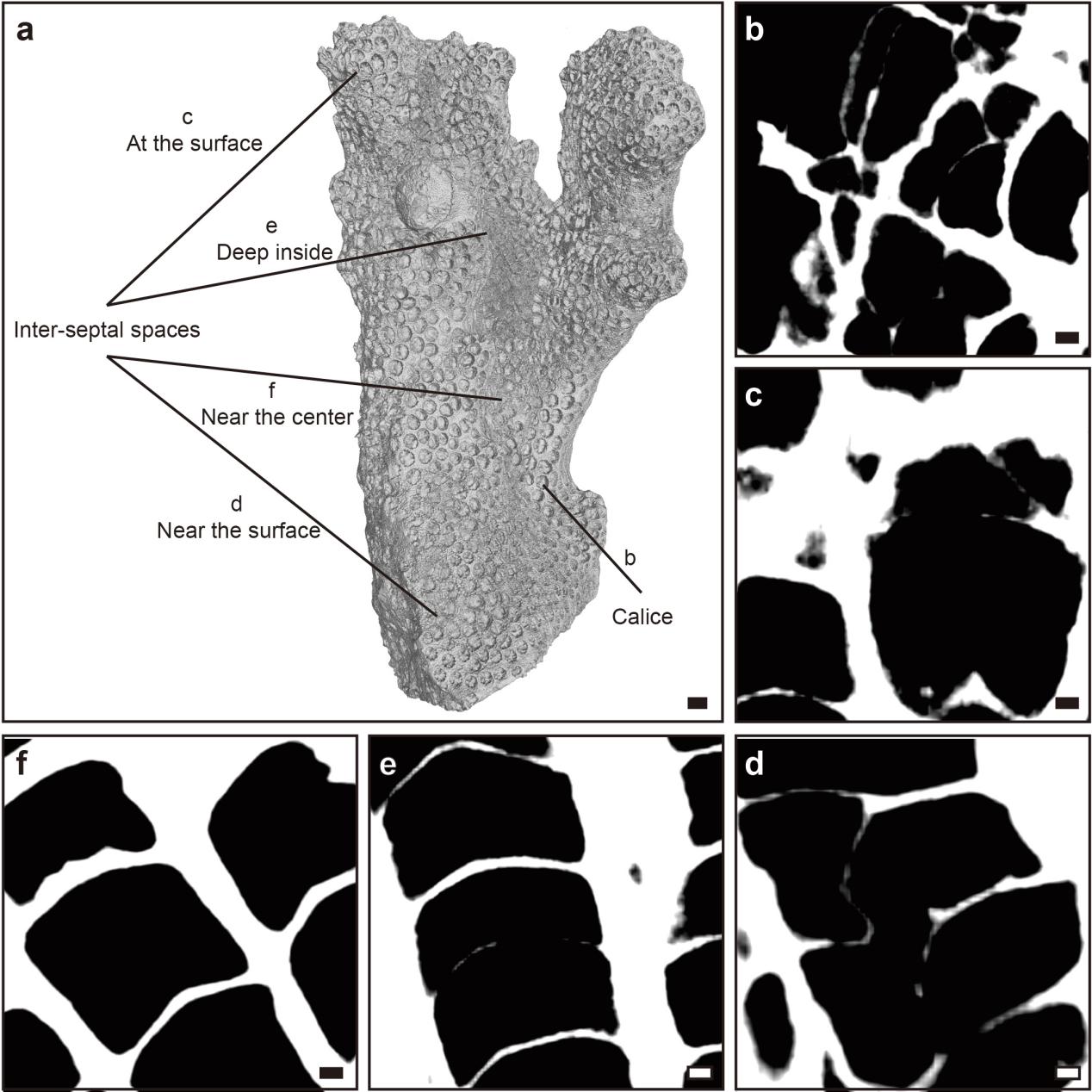

**Supplementary Figure 17 | Micro-CT reconstructions of *P. damicornis* on Day 3.** Scale bars: a) 1 mm; b-f) 0.1 mm.

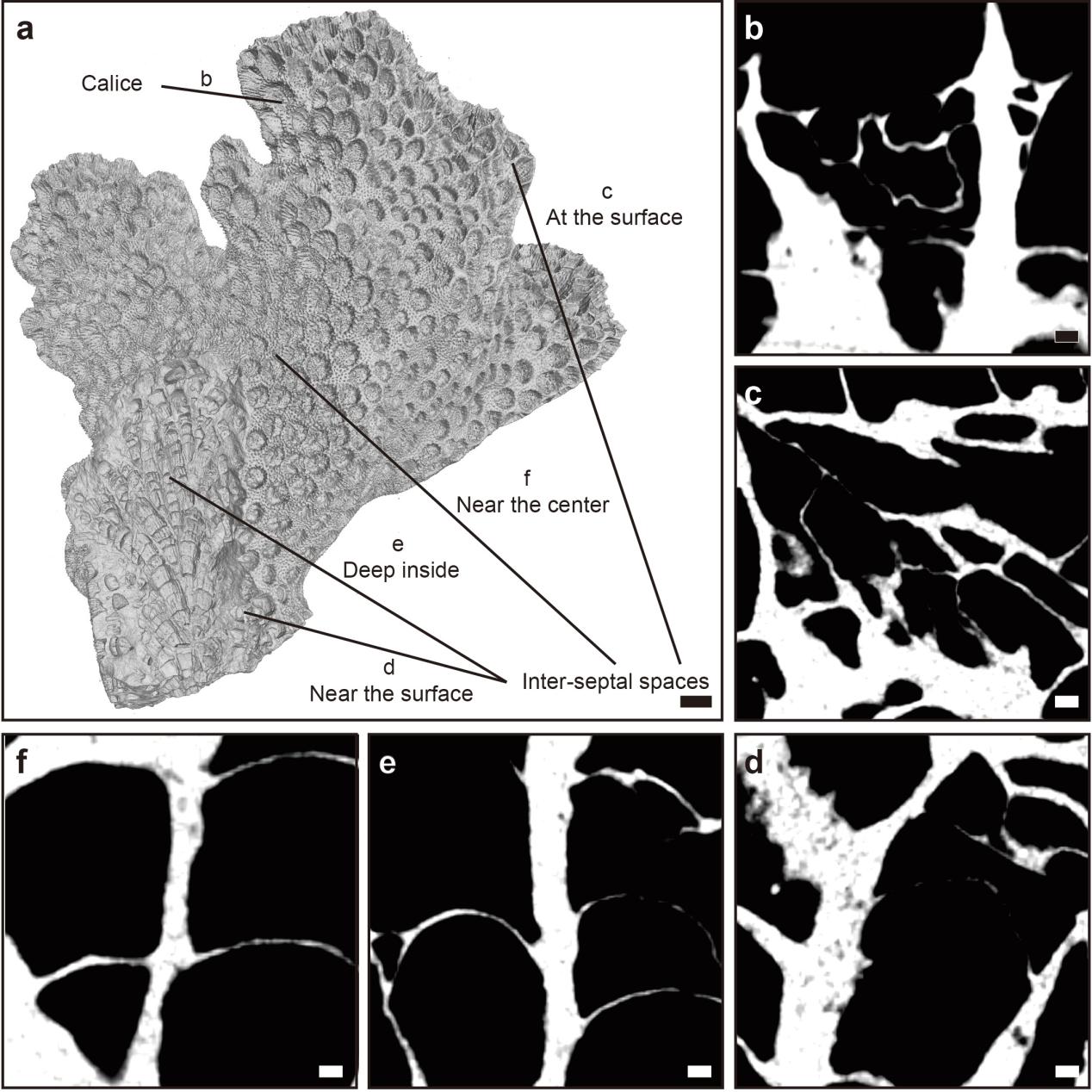

**Supplementary Figure 18 | Micro-CT reconstructions of *P. damicornis* on Day 6.** Scale bars: a) 1 mm; b-f) 0.1 mm.

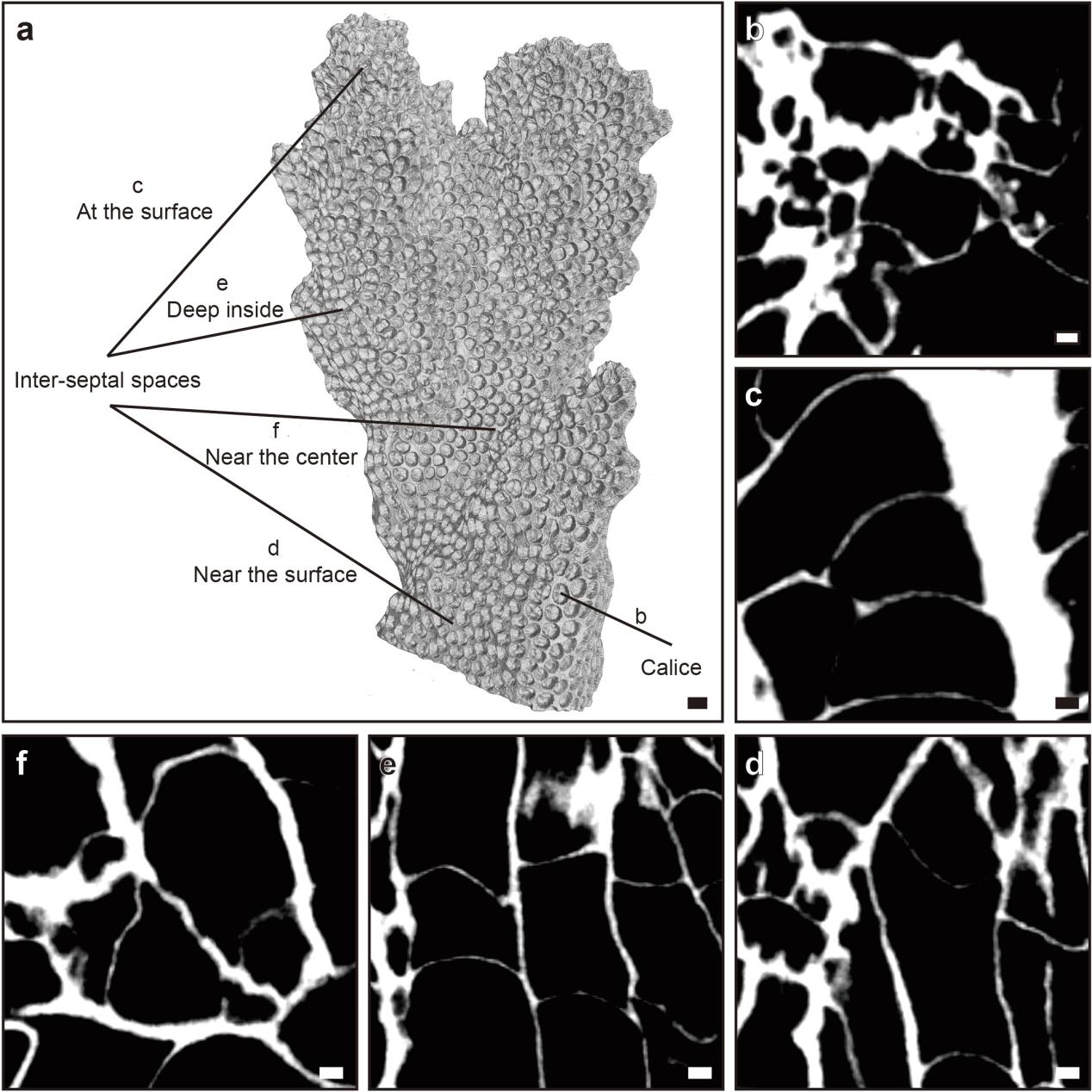

**Supplementary Figure 19 | Micro-CT reconstructions of *P. damicornis* on Day 9.** Scale bars: a) 1 mm; b-f) 0.1 mm.

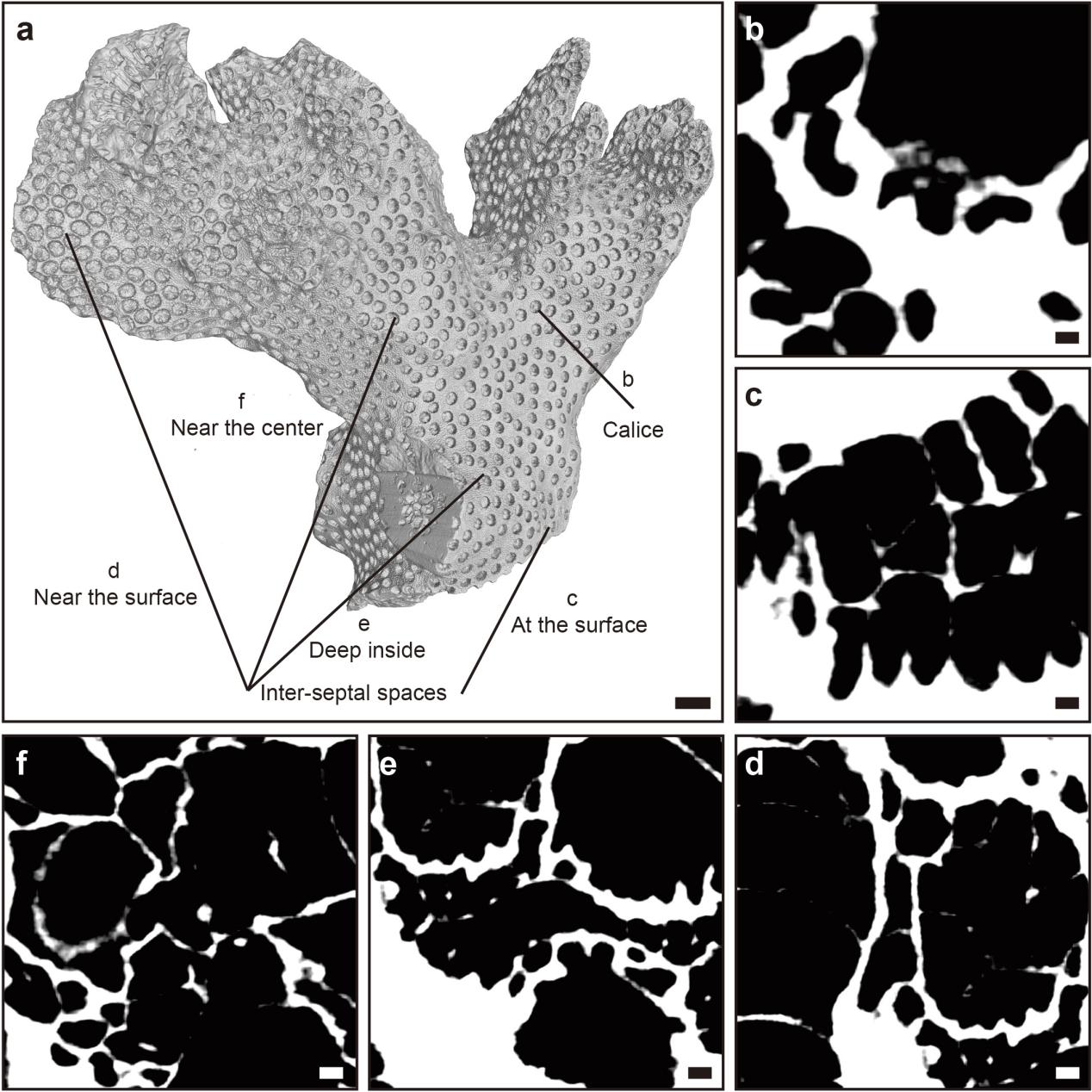

**Supplementary Figure 20 | Micro-CT reconstructions of *P. damicornis* on Day 30.** Scale bars: a) 1 mm; b-f) 0.1 mm.

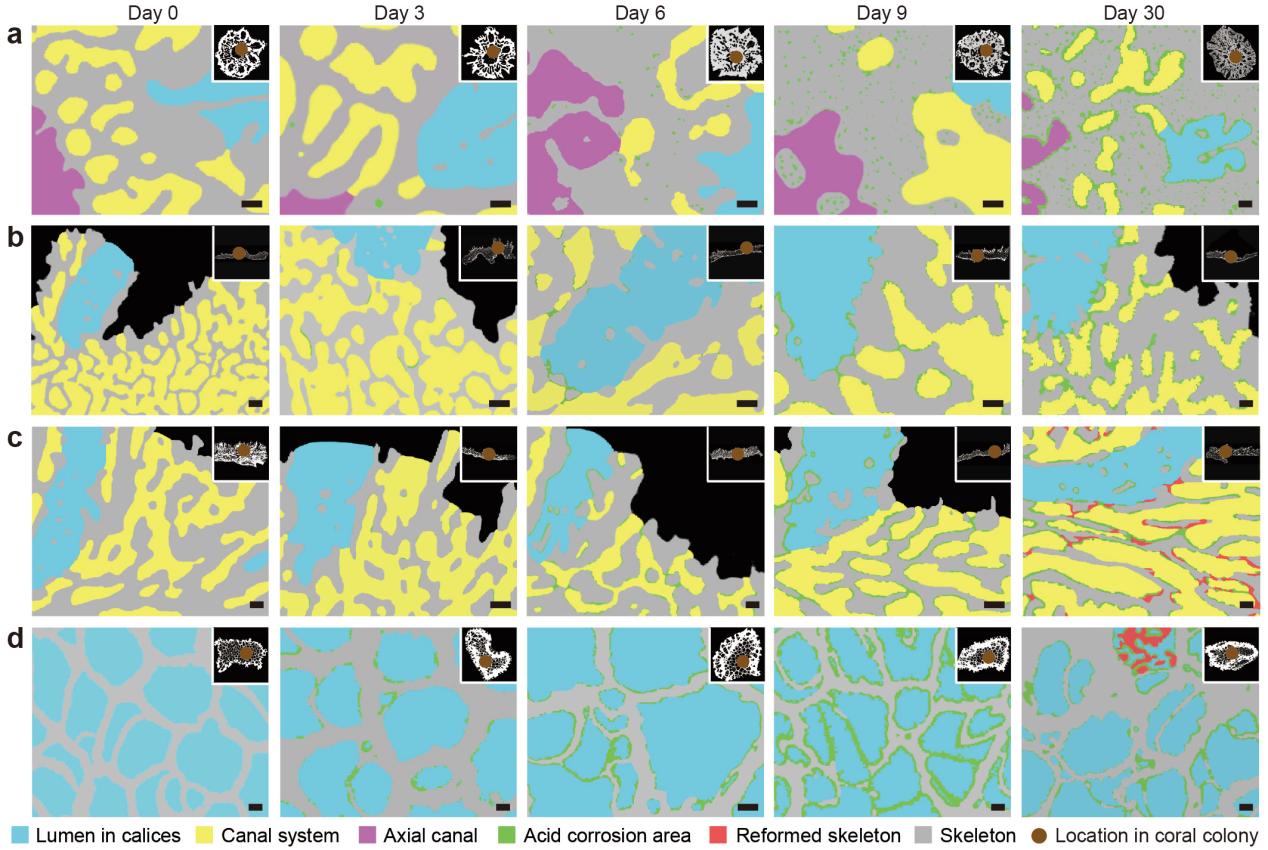

**Supplementary Figure 21 | Micro-CT reconstructions of skeletons and polyp-canal systems in coral samples from Day 0 to Day 30.** Micro-CT reconstructions visualized the erosion process in a) *A. muricata*; b) *M. capricornis*; c) *M. foliosa*; d) *P. damicornis*. Scale bars: 0.1 mm.

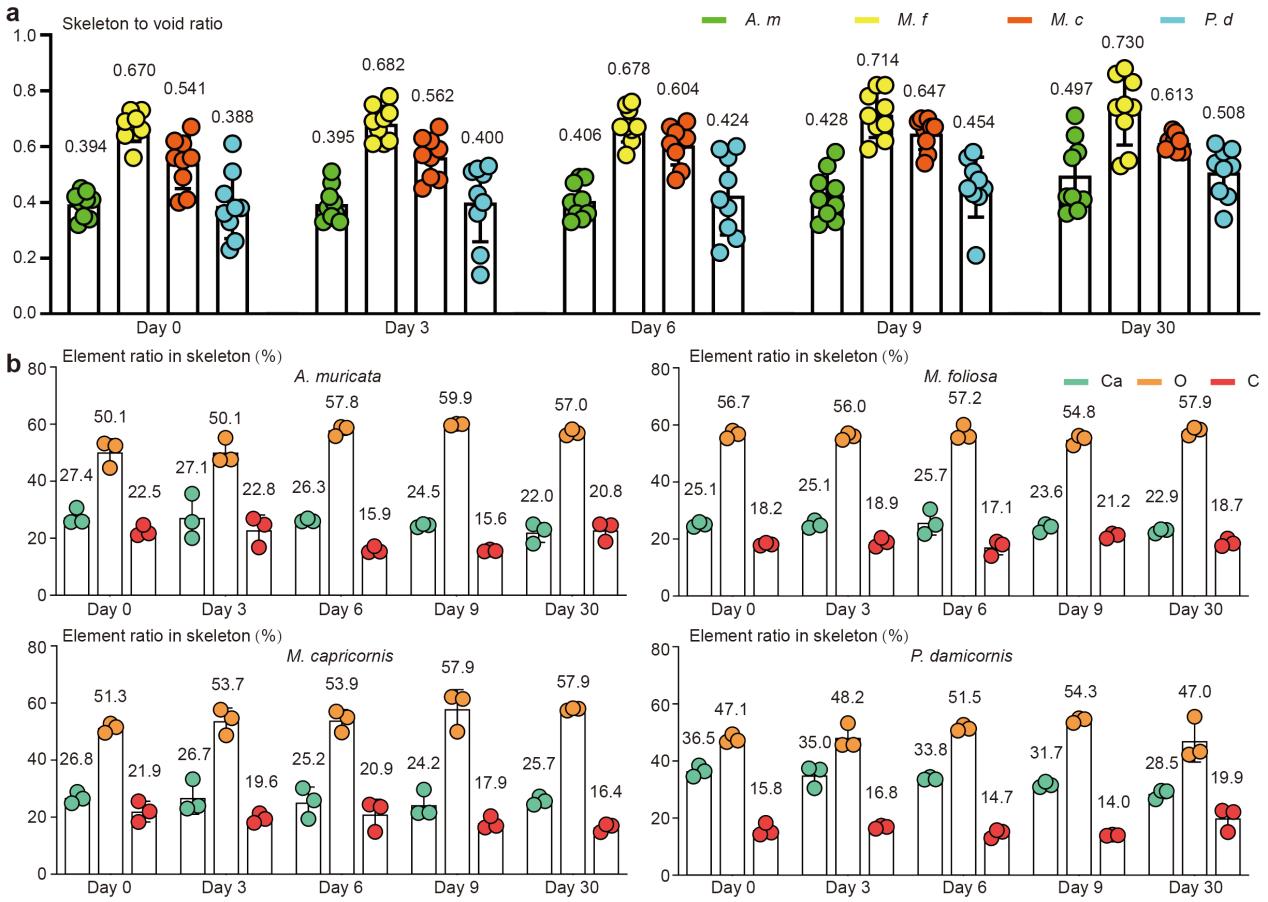

**Supplementary Figure 22 | Skeleton and element loss in coral samples from Day 0 to Day 30.** a) Skeleton to void ratio reveal the skeleton loss during acidic stress. b) Element changes of Ca, O, C reveal the affect of ocean acidification on coral skeletons.

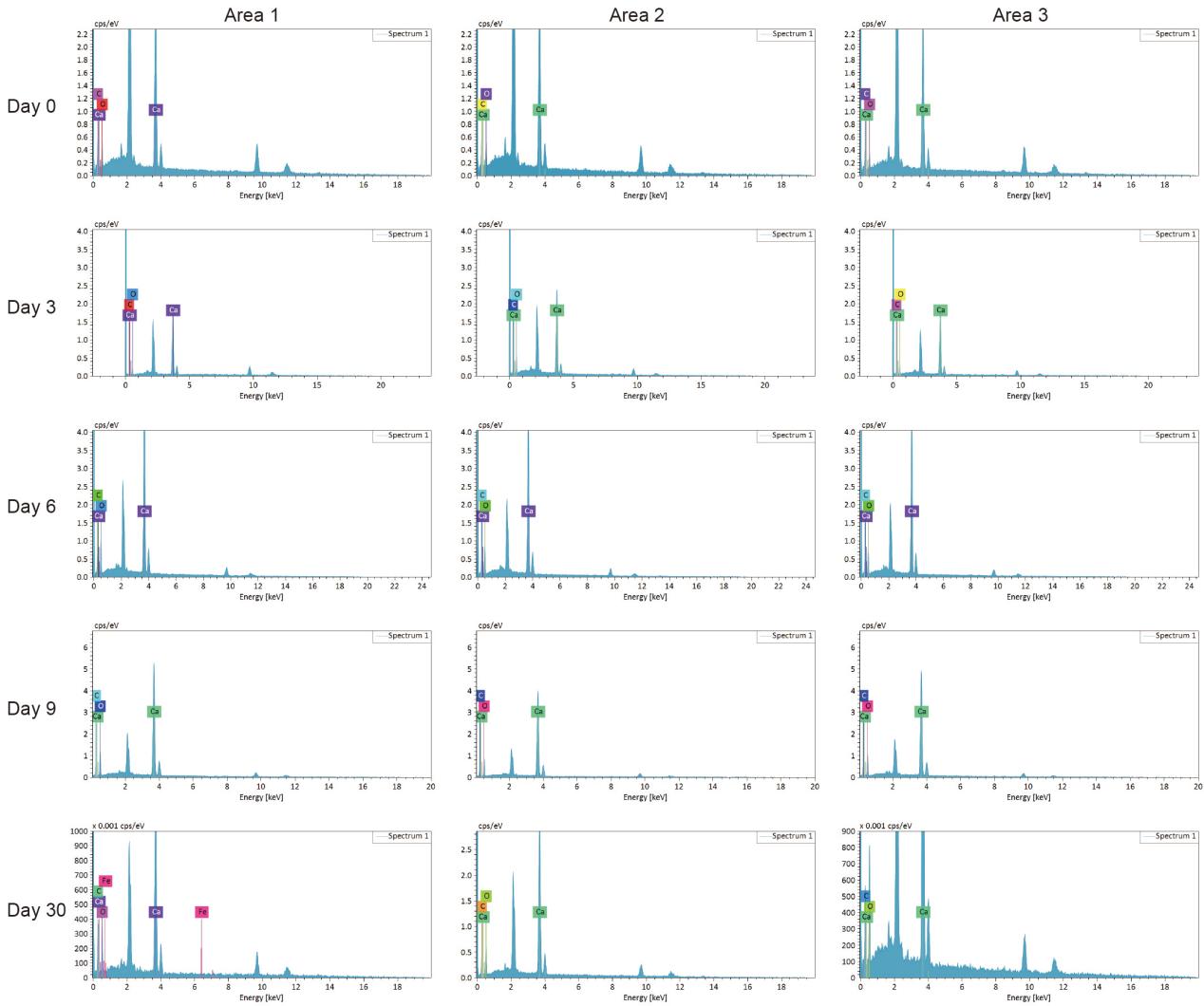

**Supplementary Figure 23 | EDS test of *A. muricata*.** The atomic ratio of Ca decreased between Days 0 and 30, with rates of decrease of 5.40% in *A. muricata*.

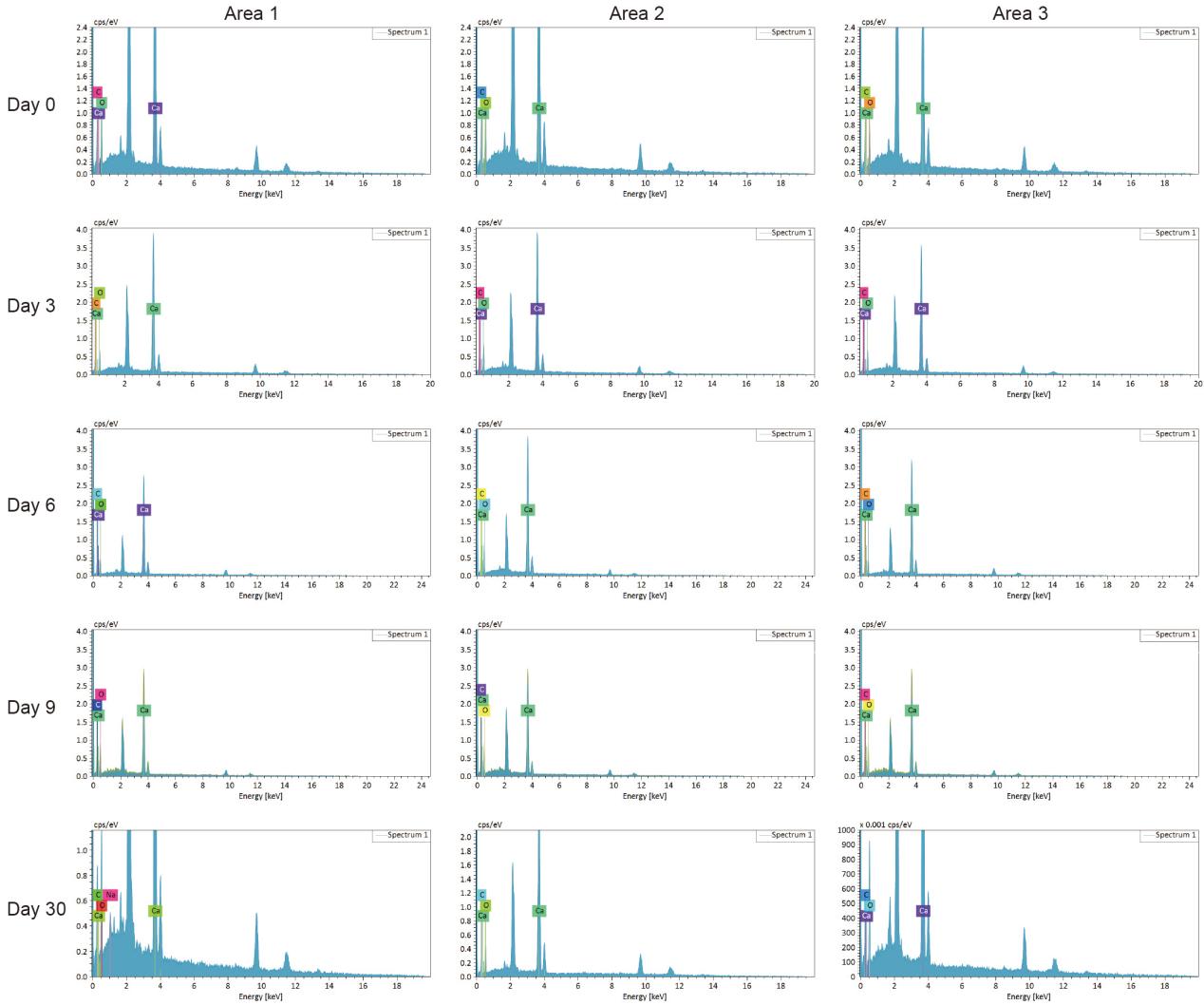

**Supplementary Figure 24 | EDS test of *M. capricornis*.** The atomic ratio of Ca decreased between Days 0 and 30, with rates of decrease of 1.01% in *M. capricornis*. The Ca atomic ratio remained constant between Days 0 and 3 in *M. capricornis*, and appeared to slightly rebound at Day 30 in *M. capricornis* from 24.24% to 25.74%.

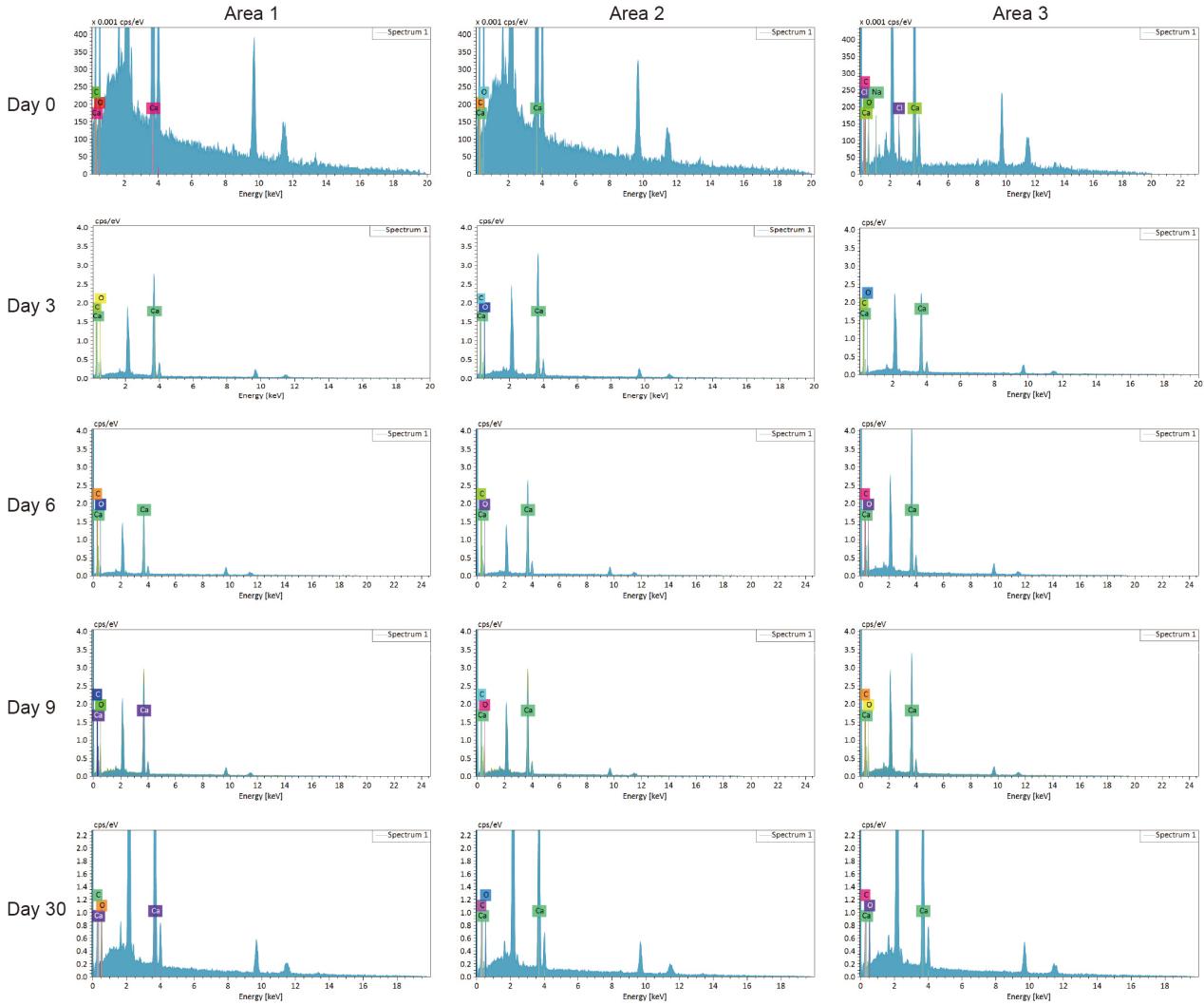

**Supplementary Figure 25 | EDS test of *M. foliosa*.** The atomic ratio of Ca decreased between Days 0 and 30, with rates of decrease of 2.26% in *M. foliosa*. The Ca atomic ratio remained constant between Days 0 and 3 in *M. foliosa*, and appeared to slightly rebound at Day 6 from 25.11% to 25.74%.

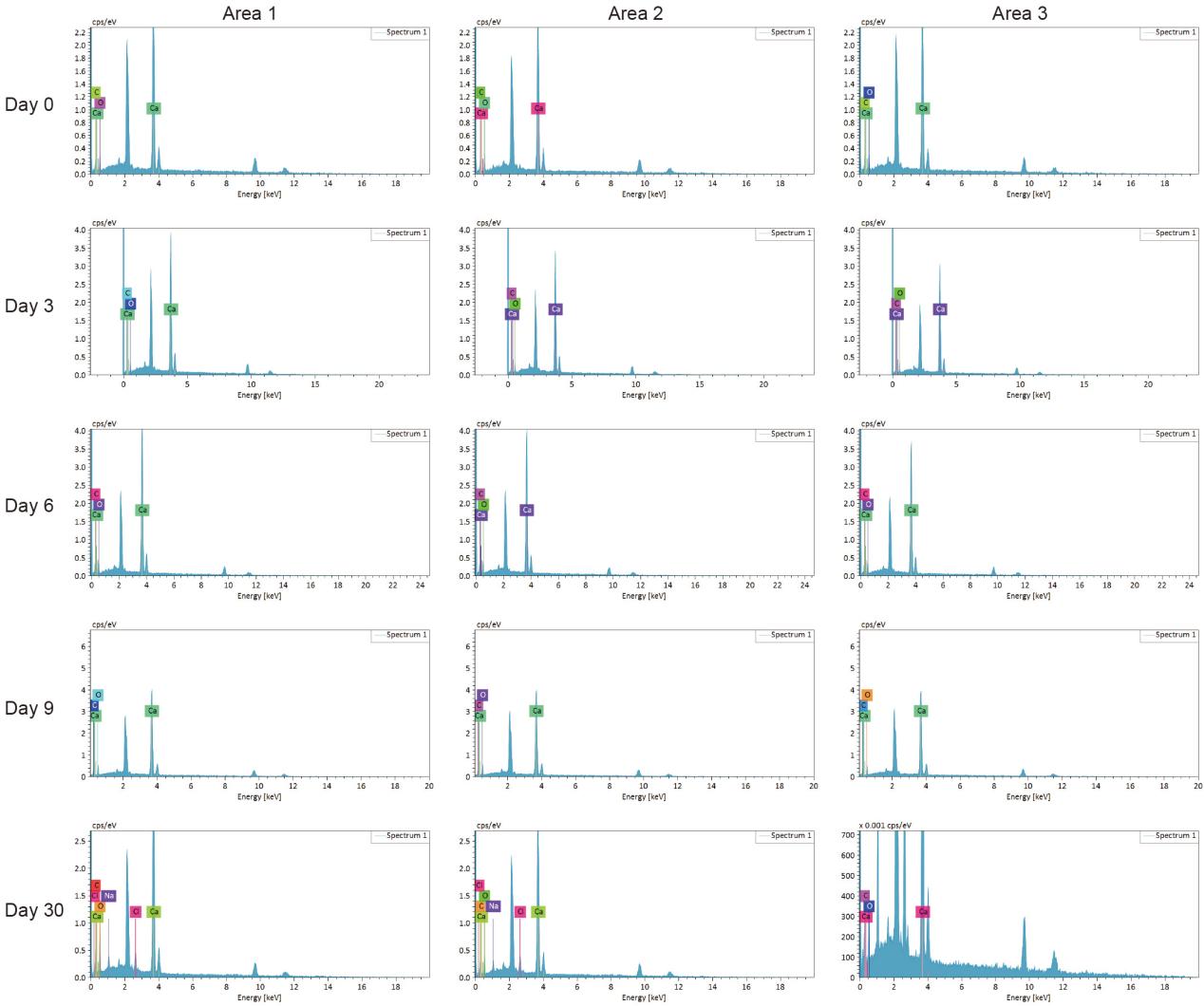

**Supplementary Figure 26 | EDS test of *P. damicornis*.** The atomic ratio of Ca decreased between Days 0 and 30, with rates of decrease of 5.94% in *P. damicornis*.

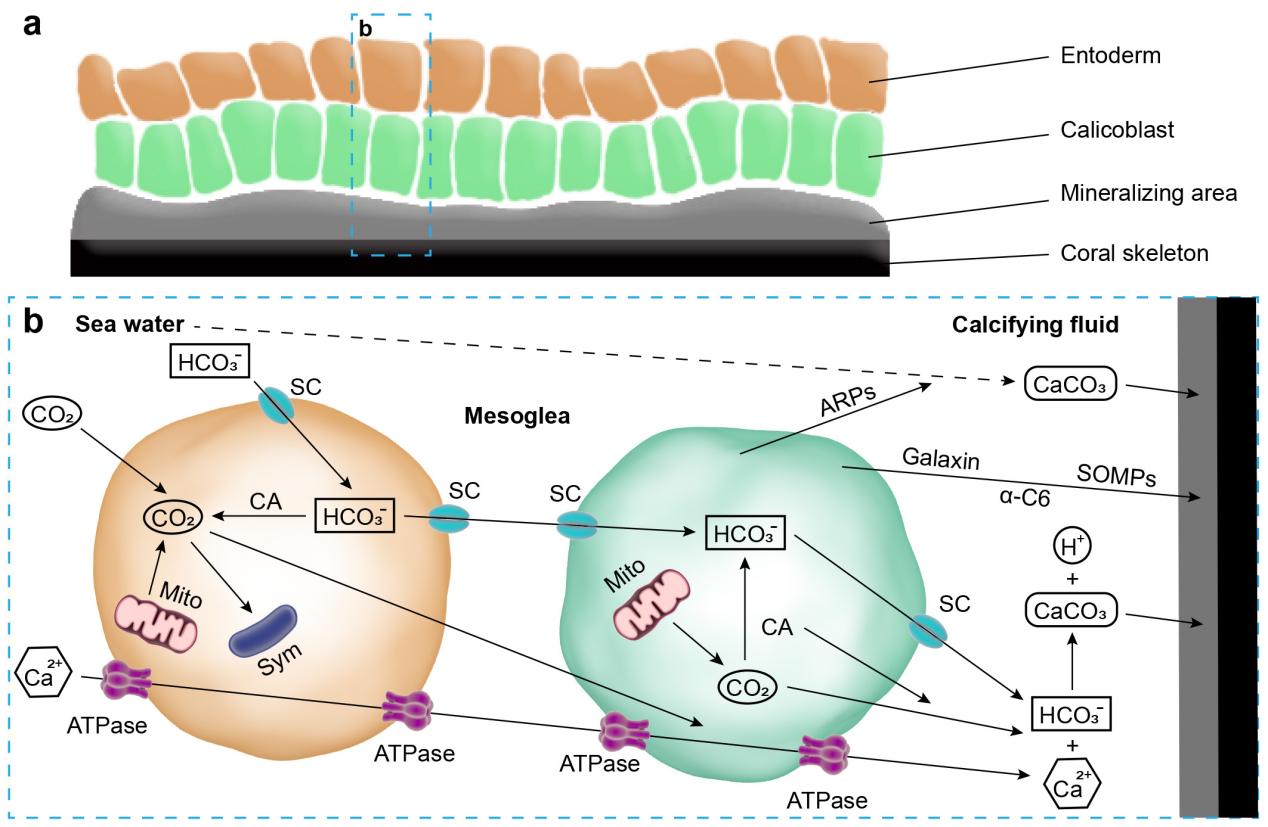

**Supplementary Figure 27 | Skeleton formation in reef-building corals.** a) Schematic diagram of coral skeletons and cells near the mineralizing area. b) ATPase means calcium ATPase; SC means solute carrier 4 and solute carrier 26; CA means carbonic anhydrase; Sym means Symbiodiniaceae; Mito means mitochondrion; ARPs means coral acid-rich proteins; α-C6 means collagen alpha-6(VI) chain-like; Galaxin means galaxin proteins; SOMPs means uncharacterized skeletal organic matrix proteins; the left cell belongs to entoderm, and the right cell belongs to calicoblast. The solid lines represent definite paths the dashed lines represent possible paths. Ca^2+^ transport by calcium ATPase or diffusion. CO_2_ can be converted into HCO^3−^ by CA and then exits the cells via bicarbonate transporters SC. ARPs can precipitate calcium carbonate from unamended seawater and modify the mineral polymorph. α-C6 and galaxin can cement the aragonite crystals to each other and to the underlying skeleton. The bioprecipitation of aragonite crystals in corals requires SOMPs.

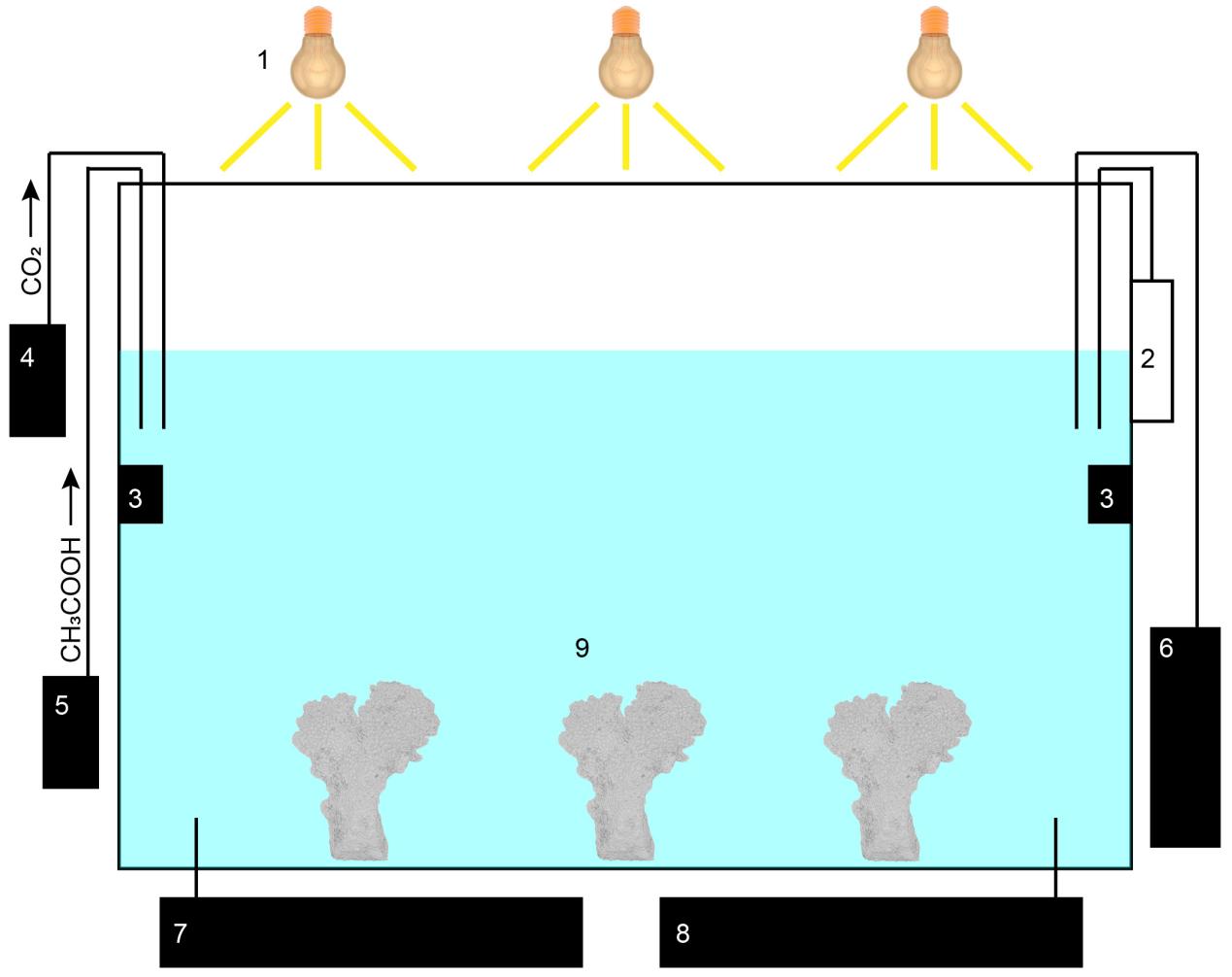

**Supplementary Figure 28 | Schematic diagram of the ocean acidification simulation device.** 1 means coral lamp, 2 means temperature and pH measuring instrument, 3 means wave device, 4 means CO_2_ bubbling device, 5 means microfluidic device for CH_3_COOH, 6 means calcium reactor, 7 means protein skimmer, 8 means water chiller, 9 means coral sample.

**Part 2 | Supplementary Tables**

**Supplementary Table 1 | Trends of skeleton to void ratio in coral colonies under acid stress.**

| **Species** | **Group** | **Day 0** | **Day 3** | **Day 6** | **Day 9** | **Day 30** |
| --- | --- | --- | --- | --- | --- | --- |
| ***Acropora***  ***muricata*** | G1 | 0.41 | 0.33 | 0.49 | 0.39 | 0.36 |
|  | G2 | 0.32 | 0.38 | 0.40 | 0.45 | 0.42 |
|  | G3 | 0.40 | 0.36 | 0.41 | 0.32 | 0.41 |
|  | G4 | 0.41 | 0.47 | 0.36 | 0.53 | 0.42 |
|  | G5 | 0.45 | 0.40 | 0.36 | 0.47 | 0.37 |
|  | G6 | 0.42 | 0.51 | 0.34 | 0.33 | 0.58 |
|  | G7 | 0.44 | 0.35 | 0.49 | 0.41 | 0.56 |
|  | G8 | 0.34 | 0.33 | 0.47 | 0.36 | 0.64 |
|  | G9 | 0.35 | 0.42 | 0.33 | 0.58 | 0.71 |
|  | Average | 0.394 | 0.395 | 0.406 | 0.428 | 0.497 |
| ***Montipora***  ***foliosa*** | G1 | 0.73 | 0.61 | 0.62 | 0.62 | 0.53 |
|  | G2 | 0.65 | 0.62 | 0.66 | 0.67 | 0.69 |
|  | G3 | 0.73 | 0.66 | 0.67 | 0.71 | 0.74 |
|  | G4 | 0.67 | 0.69 | 0.67 | 0.74 | 0.74 |
|  | G5 | 0.64 | 0.70 | 0.73 | 0.68 | 0.75 |
|  | G6 | 0.69 | 0.72 | 0.75 | 0.59 | 0.83 |
|  | G7 | 0.66 | 0.75 | 0.76 | 0.78 | 0.86 |
|  | G8 | 0.56 | 0.61 | 0.67 | 0.82 | 0.88 |
|  | G9 | 0.70 | 0.78 | 0.57 | 0.82 | 0.55 |
|  | Average | 0.670 | 0.682 | 0.678 | 0.714 | 0.730 |
| ***Montipora***  ***capricornis*** | G1 | 0.56 | 0.56 | 0.51 | 0.57 | 0.60 |
|  | G2 | 0.40 | 0.57 | 0.62 | 0.66 | 0.62 |
|  | G3 | 0.49 | 0.59 | 0.63 | 0.69 | 0.62 |
|  | G4 | 0.55 | 0.62 | 0.67 | 0.70 | 0.58 |
|  | G5 | 0.61 | 0.48 | 0.69 | 0.70 | 0.66 |
|  | G6 | 0.62 | 0.67 | 0.65 | 0.67 | 0.58 |
|  | G7 | 0.67 | 0.63 | 0.61 | 0.62 | 0.59 |
|  | G8 | 0.41 | 0.45 | 0.58 | 0.67 | 0.65 |
|  | G9 | 0.56 | 0.49 | 0.48 | 0.54 | 0.62 |
|  | Average | 0.541 | 0.562 | 0.604 | 0.647 | 0.613 |
| ***Pocillopora damicornis*** | G1 | 0.23 | 0.50 | 0.38 | 0.21 | 0.54 |
|  | G2 | 0.33 | 0.51 | 0.56 | 0.45 | 0.42 |
|  | G3 | 0.36 | 0.52 | 0.59 | 0.56 | 0.44 |
|  | G4 | 0.43 | 0.53 | 0.41 | 0.51 | 0.34 |
|  | G5 | 0.51 | 0.36 | 0.27 | 0.42 | 0.58 |
|  | G6 | 0.26 | 0.43 | 0.22 | 0.43 | 0.59 |
|  | G7 | 0.38 | 0.21 | 0.31 | 0.48 | 0.53 |
|  | G8 | 0.38 | 0.40 | 0.60 | 0.58 | 0.61 |
|  | G9 | 0.61 | 0.14 | 0.48 | 0.45 | 0.52 |
|  | Average | 0.388 | 0.400 | 0.424 | 0.454 | 0.508 |

**Supplementary Table 2 | Trends of element ratio in coral skeletons under acid stress.**

| **Species** | **Group** | **Element** | **Day 0** | **Day 3** | **Day 6** | **Day 9** | **Day 30** |
| --- | --- | --- | --- | --- | --- | --- | --- |
| ***Acropora***  ***muricata*** | G1 | Ca | 25.70% | 35.58% | 25.89% | 23.85% | 18.12% |
|  |  | O | 53.23% | 47.63% | 59.00% | 60.11% | 56.85% |
|  |  | C | 21.07% | 16.79% | 15.12% | 16.04% | 24.57% |
|  | G2 | Ca | 25.83% | 20.06% | 25.94% | 24.57% | 24.87% |
|  |  | O | 52.47% | 55.19% | 58.77% | 60.04% | 56.04% |
|  |  | C | 21.70% | 24.75% | 15.29% | 15.40% | 19.09% |
|  | G3 | Ca | 30.67% | 25.72% | 26.96% | 24.95% | 23.01% |
|  |  | O | 44.65% | 47.41% | 55.81% | 59.51% | 58.19% |
|  |  | C | 24.68% | 26.88% | 17.23% | 15.54% | 18.81% |
|  | Average | Ca | 27.40% | 27.12% | 26.26% | 24.46% | 22.00% |
|  |  | O | 50.12% | 50.08% | 57.86% | 59.89% | 57.03% |
|  |  | C | 22.48% | 22.81% | 15.88% | 15.66% | 20.82% |
| ***Montipora***  ***foliosa*** | G1 | Ca | 24.28% | 26.57% | 25.00% | 25.22% | 21.94% |
|  |  | O | 58.01% | 56.07% | 55.89% | 52.85% | 56.32% |
|  |  | C | 17.71% | 17.36% | 19.11% | 21.93% | 20.16% |
|  | G2 | Ca | 25.09% | 23.95% | 21.81% | 22.34% | 23.20% |
|  |  | O | 56.81% | 57.12% | 60.08% | 56.23% | 58.36% |
|  |  | C | 18.10% | 18.93% | 18.11% | 21.43% | 18.44% |
|  | G3 | Ca | 25.97% | 24.80% | 30.42% | 24.36% | 23.42% |
|  |  | O | 55.31% | 54.78% | 55.49% | 55.43% | 59.04% |
|  |  | C | 18.71% | 20.42% | 14.10% | 20.21% | 17.54% |
|  | Average | Ca | 25.11% | 25.11% | 25.74% | 23.64% | 22.85% |
|  |  | O | 56.71% | 55.99% | 57.15% | 54.84% | 57.91% |
|  |  | C | 18.17% | 18.90% | 17.11% | 21.19% | 18.71% |
| ***Montipora***  ***capricomis*** | G1 | Ca | 28.88% | 23.94% | 25.90% | 21.45% | 27.24% |
|  |  | O | 52.80% | 54.69% | 49.68% | 62.16% | 58.02% |
|  |  | C | 18.32% | 21.37% | 24.42% | 16.38% | 14.74% |
|  | G2 | Ca | 26.47% | 23.00% | 30.16% | 21.59% | 25.61% |
|  |  | O | 51.56% | 57.64% | 55.01% | 61.44% | 57.37% |
|  |  | C | 21.98% | 19.37% | 14.83% | 16.96% | 17.02% |
|  | G3 | Ca | 24.89% | 33.32% | 19.38% | 29.68% | 24.37% |
|  |  | O | 49.56% | 48.67% | 57.01% | 49.97% | 58.17% |
|  |  | C | 25.55% | 18.01% | 23.61% | 20.35% | 17.46% |
|  | Average | Ca | 26.75% | 26.75% | 25.15% | 24.24% | 25.74% |
|  |  | O | 51.31% | 53.67% | 53.90% | 57.86% | 57.85% |
|  |  | C | 21.95% | 19.58% | 20.95% | 17.90% | 16.41% |
| ***Pocillopora damicornis*** | G1 | Ca | 38.45% | 36.99% | 34.42% | 30.85% | 26.56% |
|  |  | O | 46.64% | 45.65% | 52.65% | 54.93% | 42.23% |
|  |  | C | 14.91% | 17.35% | 12.94% | 14.22% | 22.61% |
|  | G2 | Ca | 36.39% | 37.43% | 33.47% | 31.53% | 29.59% |
|  |  | O | 49.34% | 45.67% | 51.30% | 54.66% | 43.26% |
|  |  | C | 14.27% | 16.90% | 15.23% | 13.81% | 22.13% |
|  | G3 | Ca | 34.54% | 30.48% | 33.57% | 32.74% | 29.41% |
|  |  | O | 47.16% | 53.25% | 50.63% | 53.32% | 55.50% |
|  |  | C | 18.30% | 16.27% | 15.79% | 13.94% | 15.09% |
|  | Average | Ca | 36.46% | 34.97% | 33.82% | 31.71% | 28.52% |
|  |  | O | 47.10% | 48.19% | 51.53% | 54.30% | 47.00% |
|  |  | C | 15.78% | 16.85% | 14.65% | 13.99% | 19.94% |

**Supplementary Table 3 | Gene expression changes of coral skeletome under acid stress.**

| **Species** | **Gene type** | **Protein** | **Day 0** | | | | **Day 3** | | | | **Day 9** | | | |
| --- | --- | --- | --- | --- | --- | --- | --- | --- | --- | --- | --- | --- | --- | --- |
|  |  |  | **G1** | **G2** | **G3** | **Ave.** | **G1** | **G2** | **G3** | **Ave.** | **G1** | **G2** | **G3** | **Ave.** |
| ***Pocillopora damicornis*** | Calcium-transporting ATPase | PMC-t ATPase 2 | 177 | 313 | 363 | 284 | 86 | 86 | 92 | 88 | 35 | 14 | 44 | 31 |
|  |  | PMC ATPase | 323 | 418 | 389 | 377 | 153 | 129 | 259 | 180 | 94 | 98 | 177 | 123 |
|  | Carbonic anhydrase | CA-1 | 173 | 226 | 187 | 195 | 393 | 372 | 483 | 416 | 300 | 437 | 351 | 363 |
|  |  | CA-2 | 90 | 145 | 143 | 126 | 141 | 163 | 188 | 164 | 81 | 62 | 54 | 66 |
|  |  | CA-3 | 5913 | 8226 | 7513 | 7217 | 10374 | 10210 | 12543 | 11042 | 4170 | 5358 | 4695 | 4741 |
|  | Acidic protein | SAARP-1 | 6382 | 8179 | 7618 | 7393 | 4877 | 5012 | 5890 | 5260 | 1588 | 2040 | 1788 | 1805 |
|  |  | SAARP-2 | 190 | 212 | 189 | 197 | 2485 | 2548 | 3088 | 2707 | 5459 | 6781 | 5660 | 5967 |
|  |  | ASOMP | 98 | 132 | 130 | 120 | 207 | 146 | 254 | 202 | 77 | 102 | 86 | 88 |
|  | skeletal organic matrix protein | USOMP-3 | 798 | 1045 | 1080 | 974 | 265 | 259 | 296 | 273 | 145 | 195 | 148 | 162 |
|  |  | USOMP-5 | 824 | 1190 | 973 | 996 | 173 | 163 | 234 | 190 | 0 | 2 | 1 | 1 |
|  |  | USOMP-7 | 302 | 418 | 389 | 370 | 357 | 355 | 446 | 386 | 66 | 105 | 86 | 86 |
|  | Binding Protein | Galaxin | 438 | 593 | 576 | 536 | 8390 | 8680 | 10324 | 9131 | 9123 | 11378 | 9598 | 10033 |
| ***Acropora muricata*** | Calcium-transporting ATPase | PMC-t ATPase 2 | 517 | 478 | 656 | 550 | 37 | 70 | 79 | 62 | 6 | 0 | 18 | 8 |
|  |  | PMC-t ATPase 3 | 141 | 156 | 150 | 149 | 86 | 78 | 88 | 84 | 56 | 83 | 39 | 60 |
|  |  | PMC ATPase | 0 | 0 | 0 | 0 | 0 | 0 | 0 | 0 | 29 | 24 | 52 | 35 |
|  | Carbonic anhydrase | CA-1 | 689 | 595 | 539 | 608 | 273 | 353 | 327 | 318 | 178 | 261 | 236 | 225 |
|  |  | CA-2 | 1517 | 1626 | 1693 | 1612 | 63 | 79 | 79 | 74 | 9 | 20 | 15 | 15 |
|  |  | CA-12 | 2402 | 2163 | 2350 | 2305 | 5009 | 5846 | 6940 | 5931 | 2610 | 4054 | 3246 | 3303 |
|  | Acidic protein | SAARP-1 | 1225 | 1080 | 1241 | 1182 | 867 | 956 | 1123 | 982 | 238 | 326 | 309 | 291 |
|  |  | SAARP-2 | 638 | 577 | 616 | 610 | 239 | 266 | 290 | 265 | 1111 | 1589 | 1260 | 1320 |
|  |  | ASOMP | 7022 | 6296 | 6864 | 6727 | 1330 | 1486 | 1715 | 1510 | 1314 | 1796 | 1460 | 1523 |
|  |  | SAP-1 | 1308 | 1146 | 1380 | 1278 | 151 | 191 | 213 | 185 | 93 | 160 | 101 | 118 |
|  |  | SAP-2 | 6864 | 6450 | 7216 | 6843 | 4405 | 5098 | 5966 | 5156 | 11973 | 18929 | 14937 | 15280 |
|  | skeletal organic matrix protein | USOMP-1 | 199 | 232 | 245 | 225 | 0 | 0 | 0 | 0 | 0 | 0 | 0 | 0 |
|  |  | USOMP-2 | 494 | 434 | 429 | 453 | 42 | 42 | 65 | 50 | 7 | 11 | 1 | 6 |
|  |  | USOMP-3 | 22 | 56 | 29 | 36 | 0 | 0 | 0 | 0 | 0 | 0 | 0 | 0 |
|  |  | USOMP-4 | 4328 | 3847 | 4033 | 4069 | 3027 | 3939 | 4574 | 3847 | 755 | 1123 | 878 | 919 |
|  |  | USOMP-5 | 284 | 254 | 285 | 274 | 54 | 70 | 74 | 66 | 6 | 38 | 20 | 21 |
|  |  | USOMP-6 | 77742 | 69677 | 78447 | 75288 | 100322 | 119931 | 142703 | 120985 | 41548 | 62678 | 49748 | 51324 |
|  |  | USOMP-7 | 1251 | 1064 | 1179 | 1165 | 689 | 886 | 1022 | 866 | 560 | 967 | 726 | 751 |
|  |  | USOMP-8 | 367 | 325 | 322 | 338 | 296 | 323 | 318 | 312 | 134 | 257 | 192 | 194 |
|  | Binding Protein | α-C6 | 279 | 238 | 310 | 276 | 408 | 391 | 649 | 483 | 357 | 698 | 550 | 535 |
|  |  | Galaxin | 1981 | 1737 | 1924 | 1881 | 2803 | 3330 | 3614 | 3249 | 64382 | 98403 | 76690 | 79825 |
| ***Montipora capricornis*** | Calcium-transporting ATPase | PMC-t ATPase 2 | 1517 | 1699 | 1795 | 1670 | 1006 | 1242 | 1157 | 1135 | 1210 | 1179 | 785 | 1058 |
|  |  | PMC-t ATPase 3 | 67 | 50 | 88 | 69 | 10 | 0 | 28 | 13 | 13 | 13 | 16 | 14 |
|  |  | PMC-t ATPase 4 | 131 | 119 | 140 | 130 | 45 | 79 | 64 | 63 | 61 | 65 | 79 | 68 |
|  |  | PMC ATPase | 1367 | 1398 | 1710 | 1492 | 305 | 452 | 473 | 410 | 610 | 594 | 429 | 544 |
|  | Solute carrier | SC-4 | 426 | 490 | 573 | 497 | 353 | 438 | 499 | 430 | 550 | 340 | 438 | 443 |
|  |  | SC-26 | 3279 | 3545 | 3582 | 3469 | 818 | 944 | 849 | 870 | 2747 | 2649 | 1985 | 2460 |
|  | Carbonic anhydrase | CA-1 | 200 | 228 | 215 | 214 | 293 | 294 | 285 | 291 | 396 | 328 | 275 | 333 |
|  |  | CA-2 | 1408 | 1561 | 1440 | 1470 | 333 | 469 | 395 | 399 | 1352 | 1348 | 1014 | 1238 |
|  |  | CA-12 | 2465 | 2753 | 2906 | 2708 | 1368 | 1599 | 1465 | 1477 | 3201 | 2758 | 1863 | 2607 |
|  | Acidic protein | SAARP-1 | 981 | 1006 | 991 | 993 | 114 | 191 | 176 | 160 | 948 | 812 | 597 | 785 |
|  |  | SAARP-2 | 500 | 498 | 518 | 505 | 949 | 1144 | 1017 | 1037 | 13342 | 12549 | 9634 | 11842 |
|  |  | ASOMP | 1857 | 1920 | 1888 | 1889 | 139 | 204 | 186 | 176 | 2216 | 1880 | 1478 | 1858 |
|  |  | SAP-1 | 1207 | 1357 | 1303 | 1289 | 572 | 636 | 624 | 611 | 836 | 764 | 538 | 712 |
|  |  | SAP-2 | 1987 | 2188 | 2118 | 2097 | 2260 | 2674 | 2410 | 2448 | 5758 | 5668 | 4267 | 5231 |
|  |  | AGARP | 3373 | 3601 | 3576 | 3517 | 3239 | 3530 | 3282 | 3350 | 2669 | 2585 | 1987 | 2414 |
|  | skeletal organic matrix protein | USOMP-2 | 3207 | 3492 | 3310 | 3337 | 196 | 256 | 172 | 208 | 995 | 797 | 660 | 817 |
|  |  | USOMP-3 | 1912 | 1882 | 1947 | 1914 | 243 | 242 | 296 | 260 | 575 | 518 | 423 | 505 |
|  |  | USOMP-5 | 122 | 126 | 128 | 125 | 230 | 297 | 242 | 256 | 1317 | 1311 | 964 | 1197 |
|  |  | USOMP-6 | 13676 | 15792 | 16076 | 15181 | 21927 | 25530 | 22440 | 23299 | 25793 | 25900 | 18981 | 23558 |
|  |  | USOMP-7 | 1238 | 1319 | 1310 | 1289 | 669 | 771 | 712 | 717 | 1647 | 1516 | 1118 | 1427 |
|  |  | USOMP-8 | 32 | 56 | 52 | 47 | 72 | 83 | 51 | 69 | 24 | 25 | 23 | 24 |
|  | Binding Protein | α-C6 | 437 | 433 | 430 | 433 | 37 | 43 | 69 | 50 | 277 | 241 | 203 | 240 |
|  |  | Galaxin | 452 | 433 | 435 | 440 | 84 | 104 | 83 | 90 | 42 | 51 | 29 | 41 |
| ***Montipora foliosa*** | Calcium-transporting ATPase | PMC-t ATPase 2 | 2025 | 1980 | 1827 | 1944 | 2024 | 1427 | 1381 | 1611 | 3156 | 2402 | 3655 | 3071 |
|  |  | PMC-t ATPase 3 | 3 | 17 | 13 | 11 | 2 | 3 | 4 | 3 | 24 | 49 | 56 | 43 |
|  |  | PMC-t ATPase 4 | 455 | 431 | 455 | 447 | 181 | 110 | 104 | 132 | 167 | 134 | 192 | 164 |
|  |  | PMC ATPase | 2726 | 2598 | 2652 | 2659 | 1939 | 1496 | 1315 | 1583 | 2877 | 2057 | 3052 | 2662 |
|  | Solute carrier | SC-4 | 409 | 415 | 369 | 398 | 459 | 321 | 332 | 371 | 2486 | 1849 | 2872 | 2402 |
|  |  | SC-26 | 500 | 468 | 398 | 455 | 731 | 536 | 508 | 592 | 988 | 713 | 1193 | 965 |
|  | Carbonic anhydrase | CA-1 | 838 | 836 | 748 | 807 | 568 | 414 | 359 | 447 | 638 | 484 | 769 | 630 |
|  |  | CA-2 | 24608 | 23301 | 21649 | 23186 | 1085 | 829 | 775 | 896 | 2061 | 1469 | 2266 | 1932 |
|  |  | CA-12 | 1673 | 1727 | 1606 | 1668 | 1323 | 961 | 977 | 1087 | 773 | 560 | 882 | 738 |
|  | Acidic protein | SAARP-1 | 732 | 706 | 713 | 717 | 1065 | 702 | 765 | 844 | 1807 | 1434 | 2018 | 1753 |
|  |  | SAARP-2 | 161 | 150 | 169 | 160 | 1276 | 807 | 811 | 965 | 829 | 676 | 937 | 814 |
|  |  | ASOMP | 258 | 283 | 296 | 279 | 85 | 42 | 50 | 59 | 60 | 48 | 95 | 68 |
|  |  | SAP-1 | 1923 | 1704 | 1739 | 1789 | 986 | 703 | 628 | 772 | 1339 | 845 | 1423 | 1202 |
|  |  | SAP-2 | 9974 | 9564 | 9037 | 9525 | 10963 | 7894 | 7184 | 8680 | 5537 | 3371 | 5406 | 4771 |
|  |  | AGARP | 2630 | 2408 | 2217 | 2418 | 4522 | 3230 | 3042 | 3598 | 3161 | 2598 | 3789 | 3183 |
|  | skeletal organic matrix protein | USOMP-2 | 477 | 495 | 473 | 482 | 25 | 15 | 23 | 21 | 115 | 95 | 113 | 108 |
|  |  | USOMP-3 | 3138 | 3058 | 2849 | 3015 | 637 | 429 | 435 | 500 | 183 | 108 | 179 | 157 |
|  |  | USOMP-5 | 265 | 280 | 225 | 257 | 750 | 529 | 500 | 593 | 735 | 582 | 848 | 722 |
|  |  | USOMP-6 | 48207 | 46137 | 39596 | 44647 | 68739 | 48864 | 46750 | 54784 | 73926 | 60864 | 92171 | 75654 |
|  |  | USOMP-7 | 28 | 22 | 27 | 26 | 76 | 37 | 40 | 51 | 112 | 73 | 123 | 103 |
|  |  | USOMP-8 | 22 | 38 | 29 | 30 | 101 | 60 | 64 | 75 | 76 | 66 | 107 | 83 |
|  | Binding Protein | α-C6 | 396 | 459 | 415 | 423 | 768 | 510 | 486 | 588 | 780 | 605 | 895 | 760 |
|  |  | Galaxin | 5171 | 5105 | 4725 | 5000 | 4999 | 3634 | 3430 | 4021 | 1571 | 1222 | 1698 | 1497 |

**Supplementary Table 4 | Parameters of the micro-CT tests.**

| **Samples** | **Voltage** | **Current** | **Voxel size** | **Timing** | **Number of images** | **Image width** | **Image height** |
| --- | --- | --- | --- | --- | --- | --- | --- |
| **Day 0 *Acropora muricata*** | 120 kV | 90 μA | 9 μm | 500 ms | 4,000 | 2,300 pixels | 4,000 pixels |
| **Day 3 *Acropora muricata*** | 160 kV | 80 μA | 7 μm | 334 ms | 6,000 | 3,000 pixels | 4,000 pixels |
| **Day 6 *Acropora muricata*** | 110 kV | 80 μA | 4 μm | 1 s | 2,000 | 3,990 pixels | 4,000 pixels |
| **Day 9 *Acropora muricata*** | 130 kV | 70 μA | 3 μm | 1 s | 2,000 | 3,000 pixels | 3,800 pixels |
| **Day 30 *Acropora muricata*** | 150 kV | 70 μA | 8 μm | 500 ms | 4,000 | 2,000 pixels | 4,000 pixels |
| **Day 0 *Montipora foliosa*** | 200 kV | 100 μA | 18 μm | 500 ms | 4,000 | 2,024 pixels | 2,024 pixels |
| **Day 3 *Montipora foliosa*** | 180 kV | 90 μA | 14 μm | 500 ms | 2,500 | 4,000 pixels | 4,000 pixels |
| **Day 6 *Montipora foliosa*** | 200 kV | 90 μA | 11 μm | 500 ms | 4,000 | 3,500 pixels | 4,000 pixels |
| **Day 9 *Montipora foliosa*** | 180 kV | 80 μA | 9 μm | 500 ms | 4,000 | 3,990 pixels | 4,000 pixels |
| **Day 30 *Montipora foliosa*** | 180 kV | 80 μA | 10 μm | 500 ms | 4,000 | 3,990 pixels | 4,000 pixels |
| **Day 0 *Montipora capricomis*** | 120 kV | 90 μA | 9 μm | 500 ms | 4,000 | 3,300 pixels | 2,500 pixels |
| **Day 3 *Montipora capricomis*** | 180 kV | 90 μA | 8 μm | 500 ms | 4,000 | 3,990 pixels | 4,000 pixels |
| **Day 6 *Montipora capricomis*** | 180 kV | 80 μA | 11 μm | 500 ms | 4,000 | 3,990 pixels | 4,000 pixels |
| **Day 9 *Montipora capricomis*** | 180 kV | 80 μA | 8 μm | 500 ms | 4,000 | 3,990 pixels | 4,000 pixels |
| **Day 30 *Montipora capricomis*** | 180 kV | 90 μA | 9 μm | 500 ms | 4,000 | 3,990 pixels | 4,000 pixels |
| **Day 0 *Pocillopora damicornis*** | 220 kV | 120 μA | 7 μm | 1 s | 2,000 | 3,990 pixels | 4,000 pixels |
| **Day 3 *Pocillopora damicornis*** | 200 kV | 90 μA | 14 μm | 500 ms | 4,000 | 2,800 pixels | 4,000 pixels |
| **Day 6 *Pocillopora damicornis*** | 190 kV | 80 μA | 9 μm | 500 ms | 4,000 | 3,400 pixels | 4,000 pixels |
| **Day 9 *Pocillopora damicornis*** | 210 kV | 90 μA | 14 μm | 500 ms | 4,000 | 3,990 pixels | 4,000 pixels |
| **Day 30 *Pocillopora damicornis*** | 190 kV | 80 μA | 13 μm | 500 ms | 4,000 | 3,990 pixels | 4,000 pixels |

**Supplementary Table 5 | Carbonate chemistry in coral tank during the process of ocean acidification simulation (14 days).**

|  | Day0 | Pre-1 | Pre-2 | Pre-3 | Pre-4 | Pre-5 | Pre-6 | Pre-7 | Pre-8 | Pre-9 | Pre-10 | Pre-11 | Pre-12 | Pre-13 | Pre-14 |
| --- | --- | --- | --- | --- | --- | --- | --- | --- | --- | --- | --- | --- | --- | --- | --- |
| pH | 8.20 | 8.10 | 8.04 | 8.01 | 7.98 | 7.95 | 7.93 | 7.90 | 7.88 | 7.86 | 7.85 | 7.84 | 7.82 | 7.81 | 7.80 |
| Ca^2+^ | 399 | 396 | 400 | 399 | 401 | 400 | 400 | 402 | 402 | 402 | 402 | 402 | 404 | 405 | 410 |
| CO_3_^2-^ | 20.0 | 16.5 | 14.6 | 13.7 | 12.9 | 12.1 | 11.6 | 10.8 | 10.4 | 9.9 | 9.7 | 9.5 | 9.1 | 8.9 | 8.7 |
| HCO_3_^-^ | 112.5 | 116.5 | 118.4 | 119.3 | 120.0 | 120.7 | 121.1 | 121.6 | 122.0 | 122.2 | 122.4 | 122.5 | 122.7 | 122.8 | 122.9 |

**Part 3 | Data Availability Statement**

The datasets (reef-building coral holobionts full-length and short-read transcriptome sequencing raw data) generated during the current study are available at the Sequence Read Archive (SRA) publicly available repository, [https://www.ncbi.nlm.nih.gov/sra/].

Three generations of full-length transcriptome raw data: SAMN16237127 : Coral_OA1_day0 RNA-Seq of Pocillopora damicornis2: polyps; SAMN16237128 : Coral_OA2_day0 RNA-Seq of Acropora muricata: polyps; SAMN16237129 : Coral_OA3_day0 RNA-Seq of Montipora capricornis: polyps; SAMN16237130 : Coral_OA4_day0 RNA-Seq of Montipora foliosa: polyps.

Three generations of full-length transcriptome annotation data: SAMN16456055 : Coral_OA1_day0_Gene_Expression RNA-Seq of Pocillopora damicornis2: polyps; SAMN16456056 : Coral_OA2_day0_Gene_Expression RNA-Seq of Acropora muricata: polyps; SAMN16456057 : Coral_OA3_day0_Gene_Expression RNA-Seq of Montipora capricornis: polyps; SAMN16456058 : Coral_OA4_day0_Gene_Expression RNA-Seq of Montipora foliosa: polyps.

Second-generation transcriptome raw data : SAMN16365802：Coral_OA_1_day0_1 Pocillopora damicornis_2; SAMN16365803：Coral_OA_1_day0_2 Pocillopora damicornis_2; SAMN16365804：Coral_OA_1_day0_3 Pocillopora damicornis_2; SAMN16365805：Coral_OA_2_day0_1 Acropora muricata: polyps; SAMN16365806：Coral_OA_2_day0_2 Acropora muricata: polyps; SAMN16365807：Coral_OA_2_day0_3 Acropora muricata: polyps; SAMN16365808：Coral_OA_3_day0_1 Montipora capricornis: polyps; SAMN16365809：Coral_OA_3_day0_2 Montipora capricornis: polyps; SAMN16365810：Coral_OA_3_day0_3 Montipora capricornis: polyps; SAMN16365811：Coral_OA_4_day0_1 Montipora foliosa: polyps; SAMN16365812：Coral_OA_4_day0_2 Montipora foliosa: polyps; SAMN16365813：Coral_OA_4_day0_3 Montipora foliosa: polyps; SAMN16237439：Coral_OA_1_day3_1 RNA-Seq of Pocillopora damicornis2: polyps; SAMN16237440：Coral_OA_1_day3_2 RNA-Seq of Pocillopora damicornis2: polyps; SAMN16237441：Coral_OA_1_day3_3 RNA-Seq of Pocillopora damicornis2: polyps; SAMN16237442：Coral_OA_2_day3_1 RNA-Seq of Acropora muricata: polyps; SAMN16237443：Coral_OA_2_day3_2 RNA-Seq of Acropora muricata: polyps; SAMN16237444：Coral_OA_2_day3_3 RNA-Seq of Acropora muricata: polyps; SAMN16237445：Coral_OA_3_day3_1 RNA-Seq of Montipora capricornis: polyps; SAMN16237446：Coral_OA_3_day3_2 RNA-Seq of Montipora capricornis: polyps; SAMN16237447：Coral_OA_3_day3_3 RNA-Seq of Montipora capricornis: polyps; SAMN16237448：Coral_OA_4_day3_1 RNA-Seq of Montipora foliosa: polyps; SAMN16237449：Coral_OA_4_day3_2 RNA-Seq of Montipora foliosa: polyps; SAMN16237450：Coral_OA_4_day3_3 RNA-Seq of Montipora foliosa: polyps; SAMN16237451：Coral_OA_1_day9_1 RNA-Seq of Pocillopora damicornis2: polyps; SAMN16237452：Coral_OA_1_day9_2 RNA-Seq of Pocillopora damicornis2: polyps; SAMN16237453：Coral_OA_1_day9_3 RNA-Seq of Pocillopora damicornis2: polyps; SAMN16237454：Coral_OA_2_day9_1 RNA-Seq of Acropora muricata: polyps; SAMN16237455：Coral_OA_2_day9_2 RNA-Seq of Acropora muricata: polyps; SAMN16237456：Coral_OA_2_day9_3 RNA-Seq of Acropora muricata: polyps; SAMN16237457：Coral_OA_3_day9_1 RNA-Seq of Montipora capricornis: polyps; SAMN16237458：Coral_OA_3_day9_2 RNA-Seq of Montipora capricornis: polyps; SAMN16237459：Coral_OA_3_day9_3 RNA-Seq of Montipora capricornis: polyps; SAMN16237460：Coral_OA_4_day9_1 RNA-Seq of Montipora foliosa: polyps; SAMN16237461：Coral_OA_4_day9_2 RNA-Seq of Montipora foliosa: polyps; SAMN16237462：Coral_OA_4_day9_3 RNA-Seq of Montipora foliosa: polyps.
